# The Benchtop mesoSPIM: a next-generation open-source light-sheet microscope for large cleared samples

**DOI:** 10.1101/2023.06.16.545256

**Authors:** Nikita Vladimirov, Fabian F. Voigt, Thomas Naert, Gabriela R. Araujo, Ruiyao Cai, Anna Maria Reuss, Shan Zhao, Patricia Schmid, Sven Hildebrand, Martina Schaettin, Dominik Groos, José María Mateos, Philipp Bethge, Taiyo Yamamoto, Valentino Aerne, Alard Roebroeck, Ali Ertürk, Adriano Aguzzi, Urs Ziegler, Esther Stoeckli, Laura Baudis, Soeren S. Lienkamp, Fritjof Helmchen

**Affiliations:** Brain Research Institute, University of Zurich, Zurich, Switzerland; University Research Priority Program (URPP) Adaptive Brain Circuits in Development and Learning (AdaBD), University of Zurich, Zurich, Switzerland; Center for Microscopy and Image Analysis (ZMB), University of Zurich, Zurich, Switzerland; Neuroscience Center Zurich, University of Zurich, Zurich, Switzerland; Institute of Anatomy and Zurich Kidney Center (ZKC), University of Zurich, Zurich, Switzerland; Department of Physics, University of Zurich, Zurich, Switzerland; Institute of Neuropathology, University Hospital Zurich, Zurich, Switzerland; Department of Quantitative Biomedicine, University of Zurich, Zurich, Switzerland; Department of Cognitive Neuroscience, Faculty of Psychology & Neuroscience, Maastricht University, Maastricht, the Netherlands; Institute for Tissue Engineering and Regenerative Medicine (iTERM), Helmholtz Center Munich, Neuherberg, Germany; Institute for Stroke and Dementia Research, Klinikum der Universität München, Ludwig-Maximilians University Munich, Munich, German; Department of Molecular Life Sciences, University of Zurich, Zurich, Switzerland

## Abstract

In 2015, we launched the mesoSPIM initiative (www.mesospim.org), an open-source project for making light-sheet microscopy of large cleared tissues more accessible. Meanwhile, the demand for imaging larger samples at higher speed and resolution has increased, requiring major improvements in the capabilities of light-sheet microscopy. Here, we introduce the next-generation mesoSPIM (“Benchtop”) with significantly increased field of view, improved resolution, higher throughput, more affordable cost and simpler assembly compared to the original version. We developed a new method for testing objectives, enabling us to select detection objectives optimal for light-sheet imaging with large-sensor sCMOS cameras. The new mesoSPIM achieves high spatial resolution (1.5 µm laterally, 3.3 µm axially) across the entire field of view, a magnification up to 20x, and supports sample sizes ranging from sub-mm up to several centimetres, while being compatible with multiple clearing techniques. The new microscope serves a broad range of applications in neuroscience, developmental biology, and even physics.

## Introduction

Tissue clearing^1^ and light-sheet imaging^2^ are both century-old techniques that have garnered significant attention in the past two decades^3,4^. The combination of tissue clearing with light-sheet microscopy^5–9^ opened up new possibilities in neuroscience^10–14^, developmental biology^15–17^, and other biomedical fields^18–20^. These techniques allow researchers to visualize the anatomical structures of entire organs and even whole animals in three dimensions (3D) with high speed, contrast, and resolution, all without the need for labor-intensive tissue sectioning. The currently available techniques allow labelling and clearing of tissue samples that span multiple centimetres in size, including the whole mouse^27,28^, as well as the entire human organs such as eyes, kidneys, and even brains^29^. On the other hand, imaging smaller samples in large quantities increasingly finds applications in pathology workflows that are shifting from 2D to 3D^30–32^. This spectrum of applications requires both high resolution *and* large field of view (FOV) — two conflicting requirements in microscopy — to speed up the acquisition, minimize stitching artifacts, and maximize the information content of the image.

The challenge of maximizing resolution while increasing the FOV is not specific to microscopy. To maximize the FOV, astrophotography^33^ and machine vision industry currently employ CMOS cameras from 25 up to 600 megapixels and sensor diagonals from 35 up to 115 mm (e.g. Teledyne Photometrics COSMOS-66). Notably, several microscopy projects developed high-resolution, large field of view systems as well. The AMATERAS^34^ system employed a 122 MP CMOS camera with 35 mm diagonal and a 2x telecentric lens, and optical resolution of 2.5 µm for wide-field imaging of rare cellular events. Another example is the custom-made Mesolens^35^ objective that provides a lateral resolution of 0.7 µm across 24 mm field (sensor side), for epi-fluorescence or confocal microscopy. Most recently, the ExA-SPIM^36^ light-sheet microscope achieved an unprecedented FOV with a machine vision lens and a 151 MP CMOS camera (sensor diagonal 66 mm), with lateral and axial resolution of 1 µm and 2.5 µm, respectively.

The trend towards larger sensors in microscopy continues, and several sCMOS cameras are now available with programmable rolling shutter (‘light-sheet’) readout mode that are suitable for mesoSPIM imaging and feature large sensors (diagonals of 25-29 mm, e.g. Teledyne Photometrics Iris 15 and Kinetix, Hamamatsu Orca Lightning). However, these cameras are already incompatible with most life science microscope objectives, which are designed for smaller fields of view (18-25 mm), resulting in unsatisfactory image quality at the periphery. This issue can be resolved by using new optics that meet the standards set by modern cameras. The scientific community therefore is actively developing methods for comparing and reporting properties of microscopes and their components which are relevant to the end users^37,38^. Currently, methods to quantify the detection optics of a light-sheet microscope independently from the properties of the light-sheet excitation are rarely employed. As a result, the selection criteria of a detection objective’s performance with a large-sensor camera are often obscure.

The mesoSPIM project^21^ provides instructions for building and using a facility-grade light-sheet microscope for imaging large, cleared samples in a free and open-source way. The mesoSPIM system achieves uniform axial resolution across cm-scale FOV by using the axially swept light-sheet microscopy principle (ASLM)^21,23–25^. In brief, the axially most confined region of the light-sheet (the excitation beam waist) is moved through the sample in synchrony with the camera’s programmable rolling shutter by using an electrically tunable lens (ETL) as remote focusing device. The synchrony of the beam translation and the camera readout leads to uniform axial resolution across the field of view (see reviews in Refs^25,26^). Our previous mesoSPIM design (v.5) has several limitations. It’s detection path relies on a macro zoom microscope (Olympus MVX-10), which not only limits system’s magnification range, but also makes it incompatible with the newest generation of large-sensor cameras, due to vignetting at the periphery of the sensor. The footprint of mesoSPIM v.5 is relatively large, with multiple custom parts, and requires an optical air table, thus making the microscope relatively expensive, complex to build and align for non-experts, and difficult to move to new places.

Here, we present a new version of the mesoSPIM (“Benchtop”) that features a large-sensor sCMOS camera and an optimized detection system, resulting in higher resolution across a much larger field of view. We achieved this by developing a new method to quantify the optical properties of detection objectives independently of the light-sheet excitation. The new design is more compact, more affordable, easier to build for non-experts, and it is travel-friendly. We demonstrate examples of objective performance quantification, as well as several applications of the Benchtop mesoSPIM from neuroscience and developmental biology, as well as a novel use in physics, namely imaging particle tracks in transparent crystals that work as particle detectors. We freely share full details of Benchtop mesoSPIM assembly and operation aimed at non-experts who want to build their own system.

## Results

We are introducing the Benchtop mesoSPIM (**Fig. 1a,b,c**), which offers several advantages over its predecessor, the mesoSPIM v.5, including higher resolution, a larger field of view, and a wider range of magnifications. In addition, the Benchtop mesoSPIM significantly reduces the instrument’s footprint and cost (**Table 1**), does not require an optical table, and can be used on a lab bench. The instrument is compatible with modern large-sensor sCMOS cameras, such as the Teledyne Photometric Iris 15, and offers a 1.9x larger sensor area and 3.6x higher pixel count per image than its predecessor (**Fig. 1d**). It achieves optical resolution down to 1.5 µm laterally and 3.3 µm axially, and high field flatness across the entire field (**Fig. 1e** and **SI Figs. 1, 2**). These features allow imaging of 1 cm^3^ sample volume in as little as 13 minutes (see **SI Note 1**), making it possible to image large biological samples at high speed.

**Fig 1.**
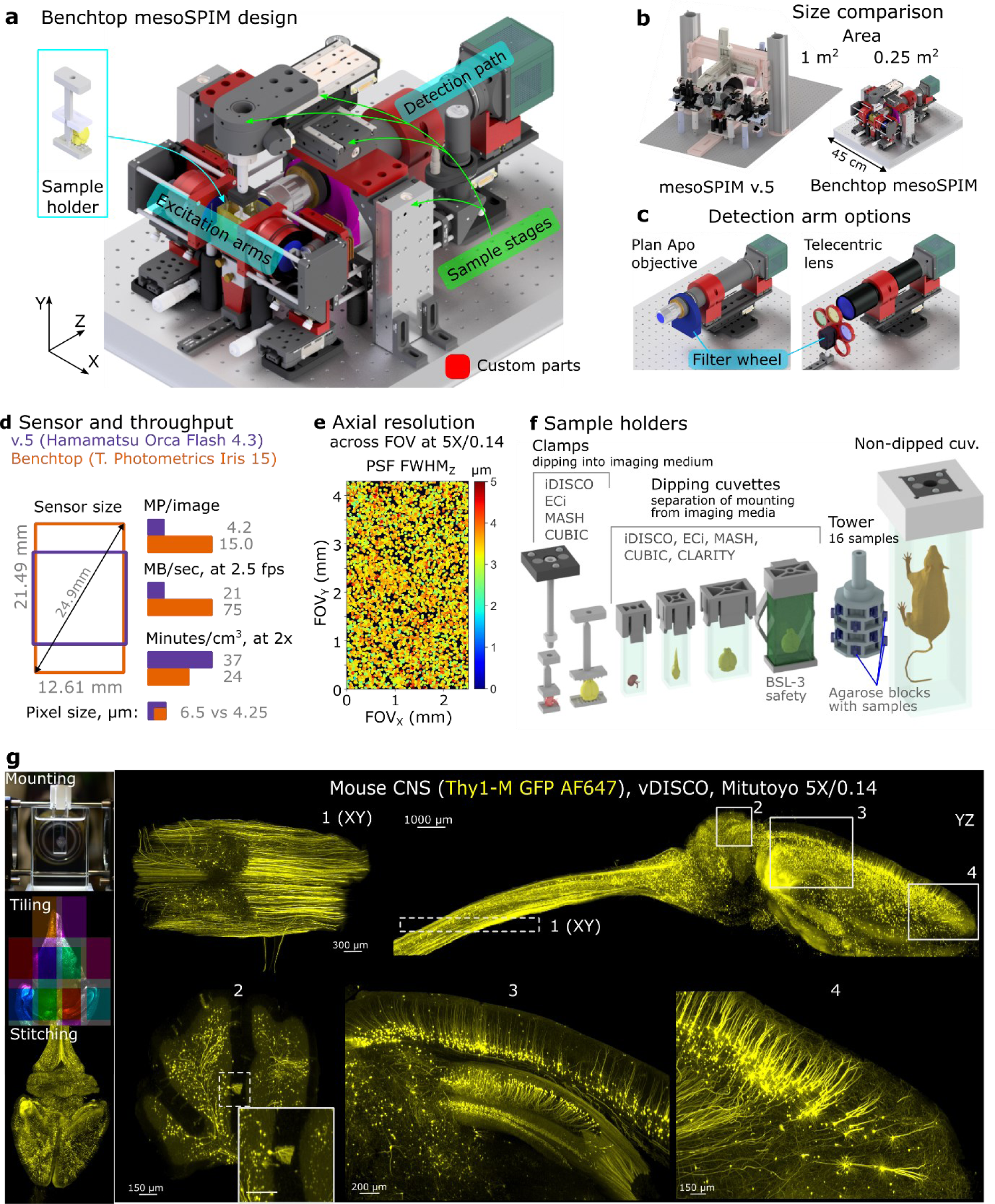
The Benchtop mesoSPIM design and an application example. **a,**CAD model of the microscope with main modules labelled: excitation arms, detection path, sample stages, and sample holder. Modified or custom-made parts are red. **b,** Size comparison between mesoSPIM v.5 and Benchtop systems. **c,** Detection arm can be equipped with a plan apochromatic objective (2x-20x magnification, tube lens not shown) or a telecentric lens (0.9x- 2x) depending on the application, with a corresponding filter wheel and set of filters. The detection arm is mounted on a focusing stage. **d,** Comparison of sensor size, pixel size, pixel count per image, and imaging throughput between v.5 (Hamamatsu Orca Flash4.3 camera) and Benchtop (Teledyne Photometrics Iris 15). **e,** The Benchtop mesoSPIM axial resolution in ASLM mode across the field of view, at magnification 5x and NA 0.14. The full width at half-maximum of point-spread function along the z-axis (FWHMZ) is color-coded from 0 to 5 µm. The resolution was measured with 0.2 µm fluorescent beads embedded in agarose immersed high-index medium (RI=1.52). **f,** CAD models of custom 3D printed sample holders that accommodate samples from 3 mm to 75 mm. **g,** Example of a multi-tile imaging of a mouse CNS (Thy1-GFP line M, Atto647N, cleared with vDISCO) at 5x magnification. The workflow consists of sample mounting, tiled acquisition, and stitching. This cm-scale sample was imaged at about 3 µm resolution that shows axonal and dendritic arborizations of long-projecting neurons: motor axons in the spinal cord (inset 1), Purkinje cells (inset 2), pyramidal cells in cortex and hippocampus (inset 3), pyramidal cells in the prefrontal cortex (inset 4). Maximum intensity projections (MIPs) over multiple planes spanning 500 µm (insets 1 and 2), 1000 µm (insets 3 and 4), or the entire volume (YZ projection of the brain) are shown. Gamma correction (1.5 in Imaris) was applied.

**Table 1:** mesoSPIM v.5 vs Benchtop comparison.

|  | Version 5 | Benchtop |
| --- | --- | --- |
| Optical resolution, axial | 5.0 μm | 3.3 μm |
| Optical resolution, lateral | 2.7 μm<sup>r1</sup> | 2.0 μm (1.5 to 2.6 μm)<sup>r2</sup> |
| Magnification range | 0.63x - 6.3x | 0.9x - 20x |
| Detection objectives | Variable (motorized zoom) | Fixed-magnification (manual exchange) |
| Camera sensor diagonal (equiv. to FOV at 1x magnification) | 19 mm | 25 mm |
| Pixel size | 6.5 μm | 4.25 μm |
| Pixels/image | 4 MP | 15 MP |
| Footprint | 1 m<sup>2</sup> | 0.25 m<sup>2</sup> |
| Mobile | no | yes |
| Approximate parts cost, USD | 162,000<sup>c1</sup> | 95,000<sup>c2</sup> |

r1, resolution estimated with Olympus MV MPLAPO 1x objective at 4x zoom<sup>21</sup>.
r2, resolution averaged across three Mitutoyo BD Plan objectives: 5x: 2.6 μm, 10x, 1.8 μm, 20x, 1.5 μm, see **SI Fig 1** for details.
c1, Version 5 costs include 4 laser lines (405, 488, 561, 638 nm) and optical table
c2, Benchtop cost estimate includes 3 laser lines (488, 561, 638 nm). Forth laser line (e.g. 405 nm) is optional, at additional cost of about 6,000 EUR. Magnifications included in the price: 2x, 5x, 10x.
Cost estimate does not include PC workstation and a screen.

To increase the system throughput and simplify sample mounting, we designed a range of sample and cuvette holders that can be 3D printed from chemically resistant plastics (polyamide PA12, see **Fig. 1f**). These holders are compatible with various clearing and immersion chemicals such as benzyl alcohol/benzyl benzoate (BABB), dibenzyl ether (DBE), and ethyl cinnamate (ECi). They are also easy to clean and autoclave. The holders can accommodate samples ranging from 3 to 75 mm in length. There are four classes of holders available (from left to right in **Fig. 1f**): clamps for direct sample dipping, holders for glass cuvettes that separate the immersion and imaging media (including extra-safe holders for BSL-3 biosafety level), high-throughput multi-sample holders (the “SPIM-tower”), and holders for non-dipped large cuvettes for extremely large samples, such as whole mice. With this diverse set of holders, our system provides high versatility for imaging a variety of samples.

Imaging the distribution of fluorescently labelled neurons in the mouse brain and their projections can be done with the Benchtop mesoSPIM at single-axon resolution when using sparsely labelled neuronal populations. A Thy1-GFP line M^39^ expressing mouse brain with spinal cord, cleared with vDISCO^27^ shows that axons and dendrites are detectable across the brain at 5x magnification, as shown in **Fig. 1g**.

Like its predecessor, the Benchtop mesoSPIM remains compatible with a wide range of clearing methods due to the use of low-NA long-working distance (WD) objectives. It is capable of imaging large clear samples due to its long-range translation stages with 50×50×100 mm travel, covering up to 250 cm^3^ volume.

Based on the configurations outlined here, users can choose a combination of detection objective, tube lens, and camera that provides a system resolution tailored to their needs. An informed choice of these fixed optical components is thus critical for good system performance.

### Testing of microscope objective contrast properties

Quality control of microscope objectives by the manufacturers is typically done manually using interferometric methods, which requires special equipment and skills. The microscopy community developed its own testing methods that aim to estimate resolution, field flatness, and other parameters across the imaging field by mounting fluorescent beads on slides (or embedding them in agarose), or by using commercially available calibration structures^38,40^.

In light-sheet microscopy, imaging of fluorescent beads embedded in agarose and index-matched with the immersion medium and measuring the point-spread function (PSF) size is currently the gold standard^25^. However, the results of such a test depend on both excitation and detection optics since the properties of the illumination beam are critical for the system’s axial resolution. In light-sheet microscopes that use ASLM scanning, such as mesoSPIM and ctASLM^24^, resolution measurements are also affected by the synchronization of the rolling shutter with the axial motion of the beam, by the exposure time, and by the lateral scanning frequency of the beam^25^. As a result, it is difficult to separate the detection objective properties from excitation parameters by measuring the system PSF from images of fluorescent beads.

To address this problem, we drew inspiration from modern methods of full-field camera lens testing^41,42^ to evaluate the modulation transfer function (MTF) across the full camera field and full focus range, resulting in 3D MTF graphs measured at specific spatial frequencies (e.g., 40 lp/mm, line pairs per mm) and angular orientations. To test microscope objectives, we adapted this method by using a high-density Ronchi grating slide at a horizontal line orientation. The slide was uniformly illuminated with an incoherent light source (a smartphone screen with an even white field), and a focus stack was captured with a 10-µm step size between planes.

The air objectives were tested in two conditions: first, with the Ronchi grating in air, and then with the Ronchi grating immersed in oil with a refractive index of 1.52, approximately 20 mm thick between the objective and the sample (as shown in **Fig. 2a**). This oil immersion mimics the conditions of imaging cleared tissue in a mesoSPIM setting as it introduces spherical aberration from the immersion medium and cleared tissue that are typically present in the system. The described testing method provides quantitative 3D contrast maps along the x, y, and focus dimensions of a detection objective (**Fig. 2b**, **SI Video 1**), allowing for the computation of compound performance metrics, such as field flatness (best-focus surface sag), depth of field, maximum contrast, and contrast variation across the field. This approach is low-cost, applicable to all types of objectives and immersion media, independent of light-sheet illumination, and can be completed in just a few minutes per objective.

**Fig. 2.**
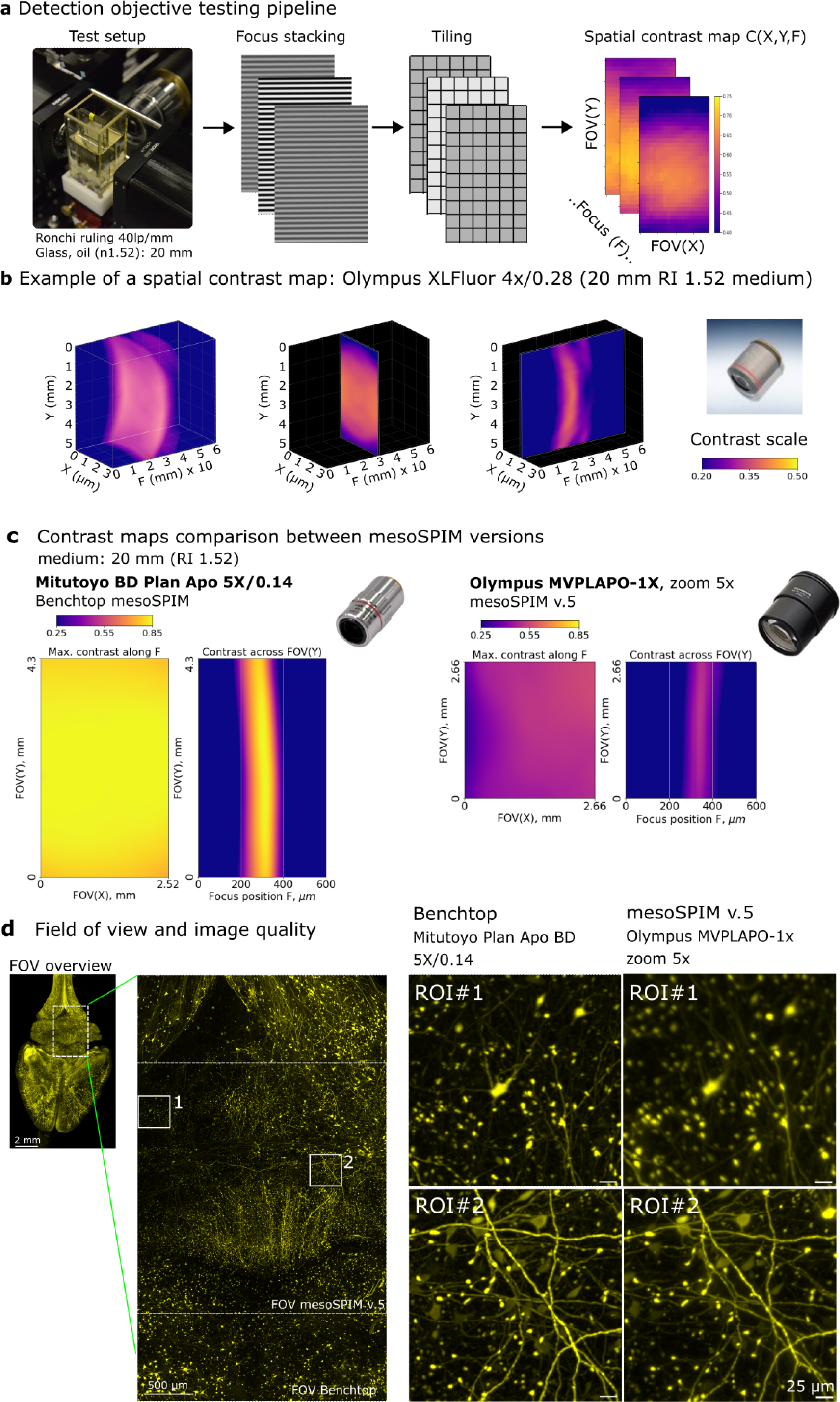
The testing method for detection objectives. **a,**The testing procedure of the detection objective contrast properties includes taking a focus stack of the Ronchi ruling immersed into imaging medium; splitting each image into sub-regions (tiling); and calculating the local contrast of each sub-region. **b,** Example of a 3D contrast map for objective Olympus XLFluor 4x/0.28. **c,** Comparison of absolute contrast and field flatness across the field between Benchtop mesoSPIM (Mitutoyo Plan Apo BD 5x/0.14) vs. mesoSPIM v5 (Olympus MVPLAPO-1x, at zoom 5x). Sensor dimensions and colormap scales are equal. **d,** Comparison of the field of view and image quality between Benchtop and mesoSPIM v.5 at 5x magnification. The Benchtop allows larger overall FOV and higher contrast across the image. The mesoSPIM v5 system has areas of lower resolution (ROI#1) and higher resolution (ROI#2) depending on the region position, consistent with the maps of objective contrast shown in **c**. MIPs over multiple z-planes are shown.

To our surprise, the contrast maps of the Olympus MVPLAPO-1x objective with zoom body MVX-10 (the default mesoSPIM v.5 configuration) showed poor centricity and large variation of contrast across field at some zoom settings, especially at 1.25x and 2x zoom (**Fig. 2c**, right panels, see further **SI Fig. 3**), presumably because of the aberrations induced by the immersion medium (see below). We thus searched for fixed-magnification objectives as an alternative, which are known to provide higher image quality at lower cost, due to the absence of moving elements and thus higher alignment precision of the lens groups. To this end we tested 11 long-WD plan apochromat air objectives from Olympus, Thorlabs, Mitutoyo, and other companies, and found their contrast performance typically degraded to various degrees at the field periphery (**SI Fig. 2,3,4** and **SI Table 3**).

**Fig. 3.**
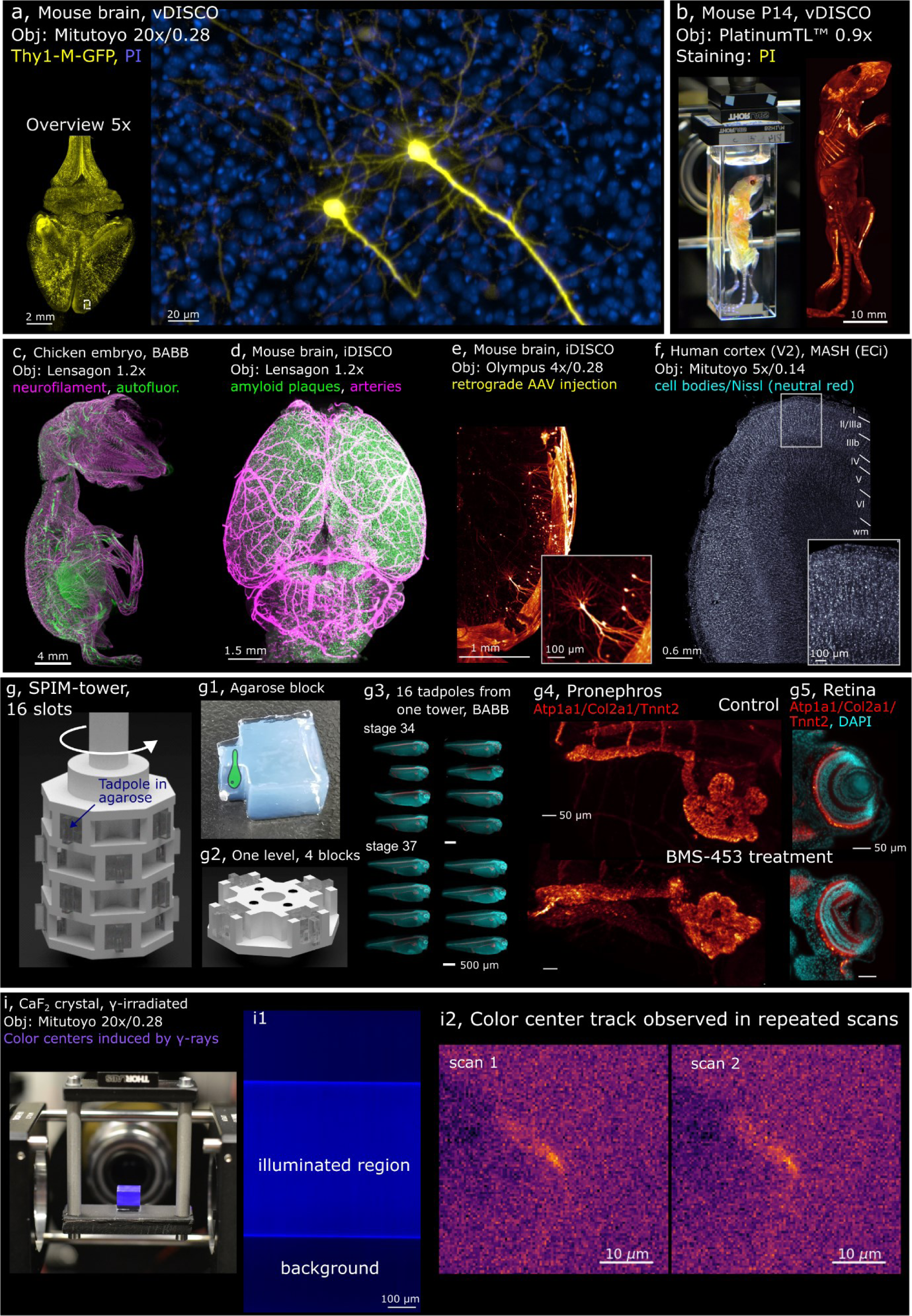
Examples of Benchtop mesoSPIM applications. **a,**Two pyramidal cells in the prefrontal cortex imaged at 20x (Thy1-GFP line M, Atto647N - yellow, propidium iodide - blue, cleared with vDISCO), with axons and basal dendrites resolved. **b,** Whole mouse body imaging at 0.9x magnification (P14 mouse, stained with propidium iodide PI, cleared with vDISCO). **c,** Peripheral nervous system of a chicken embryo at E9 imaged at 1.2x (neurofilament staining with mouse anti-RMO270, goat anti-Mouse Cy3; autofluorescence, cleared with BABB). **d,** Mouse brain at 1.2x (APPPS-1 line, amyloid plaques, arterial vessels, cleared with iDISCO). **e,** Mouse brain at 4x (Vglut2-Cre line, sparse retrograde AAV injection, iDISCO). **f,** Human brain tissue at 5x (area V2, stained with neutral red, cleared with MASH). **g,** CAD model of the *SPIM-tower* sample holder and the imaging results from high-throughput imaging session. **g1,** a molded agarose block depicting the sample as a cartoon drawing. **g2,** CAD model of a single layer of the *SPIM-tower* sample holder **g3,** Standardized imaging of 16 *X. tropicalis* tadpoles (stages 34 and 37) treated with BMS-453 (left) or control (right). Samples were stained with DAPI (cyan) and for Atp1a1/Col2a1/Tnnt2 (red), embedded in individual agarose blocks using the *SPIM-mold*; **g4**, pronephros at stage 42 (top: control, bottom: treated), **g5** retina (top: control, bottom: treated). BMS-453 treatment affects both kidney and retinal development. **i,** Irradiated CaF2 crystal imaged at 20x magnification (color centers induced by gamma irradiation at 5 MRad, polished crystal, no clearing); **i1**, raw image (color-coded blue) showing SPIM-illuminated region vs background fluorescence in the irradiated CaF2 crystal; **i2**, Candidate track of color centers observed in repeated scans of the irradiated CaF2 crystal. The structure appears across five planes (z-step 3 µm), so the maximum intensity stack is shown for each scan. In all panels, MIPs over multiple planes are shown.

We found that industrial objectives from Mitutoyo (specifically the Plan Apo BD and G series) provided the best contrast and field flatness. These objectives demonstrated excellent performance in terms of field flatness, contrast uniformity, and absolute contrast values (as shown in **SI Fig. 2**), although they are more affordable than life science microscopy objectives. When compared to the Olympus MVPLAPO-1x at the same effective magnification, the contrast maps of the Mitutoyo objectives were significantly better in terms of their absolute values, uniformity, centricity, and the resulting image quality of a biological sample (as shown in **Fig. 2c and d**). Furthermore, the availability of Mitutoyo long-WD air parfocal objectives at 2x, 5x, 7.5x, 10x, and 20x magnifications make this line optimal for a wide range of mesoSPIM applications.

Other objectives which scored high in our tests were from the Thorlabs super apochromat series and the Olympus XLFluor 4x/0.28. For applications that require lower magnification (e.g., for fast screening of mouse brains with single-cell resolution and no tiling), we tested and used industrial telecentric lenses with 0.9x and 1.2x magnification (**SI Table 1**).

The testing method described above uses white light for Rochi slide illumination, where the smartphone screen emits with red, green and blue LEDs (emission spectrum shown in **SI Fig 5a**). This restriction limits quantification of chromatic effects that would be representative for fluorescence channels. To quantify the role of chromatic effects we therefore expanded our testing method with a tungsten-halogen lamp and a set of 6 bandpass filters (spectra shown in **SI Fig. 5b**), and a Ronchi slide immersed in DBE (20 mm medium before the slide).

For a control, we imaged the Ronchi slide in the air using the Olympus MVPLAPO-1x objective at 2x zoom of MVX-10 body and found relatively flat field at all chromatic channels (**SI Fig 6a**), indicating that the objective performed nominally. This was in stark contrast with our earlier measurements when Ronchi slide was immersed in oil and illuminated with white light (**SI Fig. 3**). Indeed, we could reproduce the strongly curved field again when the slide was immersed in DBE (**SI Fig 6b**), which indicates that best-focus surface becomes curved by the presence immersion medium itself. This presumably occurs because the objective is not telecentric at low zoom, so the combination of spherical aberration, coma and astigmatism distort the best-focus field and render it non-flat. Additionally, large chromatic focal shifts are visible with the DBE-immersed Ronchi slide. Notably, focus offsets relative to the shortest wavelength (420 nm) were positive in DBE but negative in air, presumably due to DBE chromatic dispersion.

We found that under our emulated “cleared tissue” conditions, both the Olympus MVPLAPO- 1x objective and the Mitutoyo objectives have significant chromatic focal offsets (i.e. best-focus planes vary for different channels), up to 400 µm between blue (420/20 filter) and red (697/75 filter) channels (**SI Fig. 6b-e**). For example, the focal plane offset between GFP (535/22 filter) and RFP (630/69 filter) channels can be between 50 and 150 µm, positive or negative, depending on the objective. In most cases the dependence of best-focus field profile on chromatic channel was rather weak (except MVPLAPO-1x at zoom 2x), which suggests that channel-dependent focus offset is the main factor to consider for optimal imaging performance.

In practical terms, the mesoSPIM detection objective must be focused differently for each channel, which is achieved through the control software (*Focus* button group, SI Fig. 8). The amount of refocus depends on the channel, medium, and objective, and is adjusted manually for each specific set of conditions.

### Control software

The control software is an open-source Python program^22^ that is user-friendly and compatible with all versions of mesoSPIM (**SI Fig. 8**). It supports various types of hardware, such as cameras, stages, and filter wheels (**SI Tables 4, 5, 6**). The underlying PyQt5 platform allows for multi-threading to efficiently handle imaging data, while specialized libraries like npy2bdv^43^ enable fast saving of multi-tile/channel/illumination acquisitions in Fiji^44^ BigDataViewer^45^ HDF5/XML and other file formats, streamlining stitching, fusion, and visualization. This format enables datasets of 1-2 TB in size to be processed on a local workstation with modest RAM requirements (128 GB). The software is modular and allows for system upgrades by modifying a single configuration file.

### Examples of application

Imaging of a Thy1-GFP line M^39^ mouse brain with spinal cord demonstrates resolution of individual thick axons and dendrites already at 5x magnification, as shown in **Fig. 1g** and **SI Video 2** (acquisition details in **SI Table 3**). When higher magnification is required, the Benchtop system can be equipped with a 20x air objective. **Figure 3a** shows two pyramidal neurons in a mouse prefrontal cortex, with basal dendrites and axons clearly visible. Notably, the detection objective (Mitutoyo Plan Apo G 20x/0.28(t3.5)) is pre-compensated to image through 3.5 mm of glass-like medium (RI 1.52), which improves image quality by reducing spherical aberration caused by medium mismatch.

The currently available clearing protocols enable labeling and imaging of cells and their projections throughout the entire body of the mouse^27,28^. Our system is capable of imaging a whole mouse (P14 age, at 0.9x magnification, **Fig. 3b**, **SI Video 3**), potentially allowing for the creation of a whole-mouse digital atlas of cell types^28^ or tracking the spread of metastases^27^.

Development of the vertebrate nervous system, and especially axonal growth, is often studied using chicken embryos as a model^46^. As an example of imaging the nervous system *in toto*, a neurofilament-stained chicken embryo at E9 stage cleared with BABB^5^ is shown in **Fig. 3c** and **SI Video 4**.

The regional and temporal heterogeneity of amyloid-β (Aβ) plaque formation in mouse brains subject to various treatments is a promising direction in Alzheimer’s research^13^. An APP/PS1 mouse brain cleared with iDISCO^47^ is shown in **Fig 3d** (**SI Video 5**) at a magnification of 1.2x. Amyloid plaques are displayed in green, blood vessels in magenta.

The neuronal projections to specific areas of the mouse brain can be labeled by fluorescently labeled retrograde ssAAV virus stereotactic injections. As an example, we injected a Cre-dependent td-Tomato labeled construct into the lateral habenula (LHb) of a Vglut2-Cre mouse and imaged the axonal and dendritic arborizations of pyramidal cells projecting to this region (**Fig. 3e** and **SI Video 6**).

In the human brain, areas V1 and V2 are among the most thoroughly studied regions and are the only two brain areas with a macroscopically visible border between them, which is defined by the end of the stria of Gennari. This anatomical landmark is also visible in other imaging modalities such as magnetic resonance imaging (MRI), making the V1/V2 border a promising target for validating high-resolution post-mortem ultra-high field (UHF) MRI data using light-sheet microscopy. However, despite V1 and V2 being highly studied areas, reliable estimates of cell densities per mm^3^ per layer are still missing, with published estimates varying widely. High-throughput imaging of Nissl-stained samples could help to derive more reliable cell density estimates for the human brain in the future. In **Fig. 3f**, we show human cortex samples (occipital lobe samples excised around the V1/V2 border) from a 90-year-old male donor (**SI Video 9).** We also provide a dataset from a 101-year-old female donor (see **SI Video 9**). The tissues were imaged at 5x magnification, cleared, and stained with either MASH-NR or MASH- MB protocol^48^ (see **SI Fig. 9 and 10** for details).

At stage 58, *Xenopus* tadpoles enter metamorphosis, a process of drastic morphological transformations and complex physiological changes. *In toto* imaging of whole animals during this transition provides a unique opportunity to uncover the molecular mechanisms governing morphogenetic changes, such as limb development, the transition from pro- to mesonephros (the adult kidney), or tail regression. Furthermore, imaging of entire intact froglets offers a comprehensive view of organs and tissues in context that provides insights into complex interactions relevant to normal developmental processes and disease models. **SI Video 10** shows a *Xenopus* tadpole at stage 58 cleared with BABB, stained with Atp1a1 antibody that labels the nervous system projections (including the olfactory system), eyes and the developing hindlimbs of the animal at this critical developmental stage.

### High throughput imaging of biological samples

For higher throughput imaging we designed a modular “SPIM-tower” holder that accepts 4 samples per level, with multiple levels stacked upon each other connected by magnets. The samples are embedded in agarose blocks which stick out of the tower frame for obstruction-free light-sheet illumination (**Fig.3g** and **SI Fig 11**). The tower is rotated in 90-degree steps and moved vertically to bring the individual samples into position for mesoSPIM imaging. The system includes a 3D printable agarose mold that allows accurate mounting of multiple samples in identical agarose blocks that are then inserted into tower slots.

The SPIM-tower enabled us to image 16 tadpoles (4 levels stacked), embedded in individual agarose blocks and cleared with BABB, in one 20-minute session (**Fig. 3g**). Standardized automated whole-embryo imaging after experimental intervention, such as chemical treatment or genetic manipulation (e.g., CRISPR/Cas9), has the potential to facilitate screening efforts by distinguishing between intrinsic variability and genuine phenotypes. Indeed, here we demonstrate the effects of transient inhibition of retinoic acid (RA) signaling during embryonic development across *Xenopus* embryos. *In toto* analysis of embryos revealed hypoplastic disorganized pronephros with shortened tubular lengths and retina/lens hypoplasia with effects on retinal layer organization (**Fig. 3 g4,5**). Additionally, the larger size of such uniform datasets makes them ideally suited for future deep learning-based automated phenotyping initiatives^17^. This system can be used for higher-throughput imaging of embryos, organoids, and larger samples of various species.

### Imaging of color centers for particle detectors

Besides imaging biomedical samples, the high-throughput and optical sectioning capabilities of the mesoSPIM can have applications in physics and geology to study fluorescent signals from defects in transparent crystals. These fluorescence-emitting defects, so-called color centers, can be induced in crystals by irradiation^49–53^. Particles such as neutrons, ions, cosmic and gamma rays can either dislocate atoms and create vacancy defects in crystalline structures and/or provide electrons to existing defects^49–53^. The resulting system can fluoresce when excited by light. By imaging these small defects with light-sheet microscopy, it is possible to probe not only the well-known interaction of particles such as neutrons with solid-state materials, but also the flux of weakly interacting particles, such as neutrinos from nuclear reactors or dark matter candidates^54–56^. Measuring the interaction of these particles requires scanning large sample volumes^54^ (kilograms of transparent crystals). To study color center formation, we irradiated a 1cm^3^ CaF2 crystal (**Fig. 3i**) with gamma rays from a ^60^Co source which made the crystal fluorescent (**Fig. 3i1**). When the crystal was imaged at 20x magnification, several track candidates were found repeatedly in more than one scan (which minimizes the likelihood that they are noise artefacts). One such track candidate is shown in **Fig. 3i2**, with statistical analysis in **SI Note 2** and **SI Fig. 12**. While these findings are in their very early stages, the ability of the Benchtop mesoSPIM to image hundreds of cm^3^ of bulk crystals at high speed and resolution is instrumental for the detection of color centers induced by rare events in passive crystal detectors. These fluorescent structures can be produced by ions, neutrons or elusive particles, such as neutrinos and dark matter candidates^54,55^.

### Travel capability

The Benchtop mesoSPIM is made travel-friendly by grouping the elements into mechanically independent modules (2 excitation paths, detection path, sample stages), which minimized the packing and alignment requirements to ship and reassemble the system. The packed components have a volume of about 0.2 m^3^ and weight 60 kg. Unpacking and reassembling the system takes less than 1 hour (**SI Video 11**), followed by up to 1 hr of test and alignment routines.

The travel capability of Benchtop mesoSPIM enables easier access to the instrument and wider collaboration opportunities among biological labs. In the context of physics (imaging rare track events), the microscope can be more conveniently used in places of difficult access, such as in underground laboratories. The laboratories are often within mines, where the non-specific irradiation (such as cosmic rays) is minimized. For the application of nuclear reactor monitoring (discussed in **SI Note 2**), the Benchtop mesoSPIM could be transported for *in-situ* sample analysis, eliminating the requirement for a dedicated setup at each location.

### Cost reduction

The substantial cost reduction of the Benchtop mesoSPIM compared to its predecessor (95k vs 165k USD) was achieved by optimizing the costs of lasers, stages, control electronics, and detection objectives. The Benchtop uses a single laser combiner with up to 4 laser channels with a built-in fiber-switching module, which allows fast and reliable switching of the laser source between the two excitation arms with minimal power loss.

The compact footprint of the Benchtop mesoSPIM is achieved through several modifications. First, the macro-zoom system Olympus MVX-10 is replaced with fixed-magnification objectives. Additionally, the excitation path is redesigned to optimize space utilization. The use of more compact translation stages allows for efficient positioning of components within a limited area. Furthermore, a low-cost compact filter wheel, obtained from a hobby astronomy shop, is chosen as a compact alternative. These modifications collectively contribute to the reduced size and enhanced portability of the system.

### Upgrading previous versions

The older versions of mesoSPIM (v.5 and earlier) can be upgraded in their detection path to achieve the resolution and magnification range of the Benchtop mesoSPIM. The upgrade includes a large-sensor camera, Mitutoyo objectives (2x, 5x, 7.5x, 10x, 20x), the motorized compact filter wheel and the objective turret (**SI Fig. 13, SI Table 7**). The total cost of upgrade including camera is about 27k USD. All components of the excitation paths and sample stages can be re-used.

### Instructions for building and operation

Building a Benchtop mesoSPIM does not require elaborate skills in optics, mechanics, or programming. Customization of several metal components requires a basic metal workshop with milling machine to modify several off-the-shelve parts. Alternatively, metal parts can be ordered online from vendors using the design files posted on project’s website. Components for which strength and long-term dimensional stability are not critical can be printed using hobby-grade 3D printer. Several electronic parts (e.g., electro-tunable lens and galvo drivers) require modification that involves basic soldering skills.

We offer a full parts list, purchase recommendations, wiki documentation, tutorials, and videos, for experts and non-experts alike, on how to build, align and operate the Benchtop mesoSPIM.

### Current limitations

Due to our design choices, we use off-the-shelve air objectives in both excitation and detection arms for compatibility with multiple clearing protocols and quick exchange of imaging media, while keeping the system’s cost low. Thus, the optical resolution of our current design suffers from spherical aberration from the medium index mismatch. Our simulations show that both excitation and detection arms operate in a diffraction-limited regime only with air objectives of NA up to 0.15 and immersion medium thickness up to 15 mm (**SI Notes 3 and 4**). Therefore, increasing the NA of detection objectives above 0.15 achieves higher resolution only for sample (incl. immersion medium) thinner than 15 mm (a “15-15” rule). The simulated PSF size is nearly independent of the refractive index of the medium in the practically relevant range (n=1.33-1.99, **SI Table 11**).

It should be noted than “field flatness” derived from the contrast maps of objectives in our testing method should not be confused with Petzval field curvature. The field flatness profile (2D) is computed from the best-focus surface (in 3D) of the objective under testing conditions. Depending on the chosen contrast criterion, the location of the best-focus surface can be slightly different. The best-focus surface becomes more curved in the presence of planar refractive interface (cuvette wall and immersion medium) due to the presence of coma, astigmatism, and non-telecentric design of some objectives, which together bend the best-focus surface relative to the ideal conditions (air). On the contrary, the well-known Petzval field curvature does not depend on the presence of planar interfaces in the system, and it is independent of spherical aberration, coma, or astigmatism. The best-focus surface we measured in this work is similar to the concept of focal surface in optics.

## Discussion

The Benchtop mesoSPIM represents a major advancement in light-sheet microscopy for large cleared tissues, surpassing the previous mesoSPIM version in terms of optical performance, throughput, and range of possible applications. Its lower cost, ease of assembly, and travel capability makes it more affordable for research groups with a moderate budget. The modular design of the system allows for easy upgrades and maintenance, making it more future-proof than commercial systems. While some expertise is required for building and upgrading, it is within the reach of microscopy tech enthusiasts and does not require special skills.

In comparison to ExA-SPIM^36^ and NODO^32^ light-sheet microscopes, the Benchtop mesoSPIM offers lower resolution and throughput, but also lower cost and higher versatility in the choice of the detection objectives. On the other hand, compared to the descSPIM^57^, the Benchtop mesoSPIM has higher cost but also higher resolution, field of view, and throughput. This niche makes our system attractive to imaging facilities with a wide range of sample types and sizes and moderately high resolution and throughput requirements.

The objective testing method developed for this project enabled the selection of optimal detection objectives to achieve higher resolution and a larger field of view. Although this method does not provide a quantification of PSF size, as other methods that employ fluorescent beads, it does not depend on the light-sheet illumination path, making it particularly useful for light-sheet systems with complex illumination techniques, by decoupling detection properties from excitation. It can be also used for comparing objectives in other light microscopy modalities, and the resulting contrast maps are intuitive for interpretation. The method can be further improved by using several orientations of the test target, test targets of various line pairs densities to probe the system MTF at multiple spatial frequencies, and by performing more advanced computational analysis of the contrast maps to deduce the optical aberrations present in the system.

The Benchtop mesoSPIM’s axial and lateral resolution can be significantly improved by designing custom excitation and detection objectives that are corrected for spherical aberration that comes from medium mismatch. We thus encourage companies that design and produce microscopy objectives to develop long-working distance air objectives with NA 0.3 to 0.5, magnification 5x to 20x, and build-in compensation of strong spherical aberrations (an equivalent of imaging though a glass block 25-35 mm thick). The ExA-SPIM^36^ system circumvented this problem by using an industrial telecentric lens with the front beam splitter removed (an equivalent of 35 mm glass block), but this lens type is not infinity corrected, has a large footprint and a low magnification range, which limits its use in the emerging field of light-sheet imaging of cleared samples.

The Benchtop mesoSPIM can be made more compact through customized electronic components such as stage controller, waveform generator, power supplies and laser combiner. The mechanical design can be further reconfigured to accommodate an off-the-shelve motorized turret with multiple detection objectives for higher user convenience. Although our assembly and some custom parts are licenced as open-source (GPL-3), we also use multiple closed-source components, and there is a room for custom components from commercial manufacturers, if the component’s interface and performance are clearly specified and the design is not derived from already existing GPL-licenced parts.

Although we provide specific suggestions for the optomechanics, objectives, and other hardware for the Benchtop mesoSPIM, they are not exhaustive, and modifications can be made by other developers to further improve imaging quality, reduce cost, and adapt the system to new applications. We encourage open, test-based, and well-documented modifications to the system and its software, which can be shared with the imaging community through Github repositories.

We expect the Benchtop mesoSPIM and its modifications to have numerous applications in neuroscience, developmental biology, digital pathology, and physics.

## Methods

### Optomechanical design

Continuing the mesoSPIM original design goals, we opted for long working distance air objectives with low NA (< 0.3) for both detection and excitation, which ensures compatibility with all clearing methods and straightforward tiled imaging of large specimens.

The detection path can have either a *microscope* configuration (objective + tube lens) or a *telecentric lens* configuration (telecentric lens attached directly to the camera), see **Fig. 1c** and **SI Fig. 5**. Custom-made lens and camera holders (red parts) are 3D printed from PETG or similar mechanically stable plastic. The objective-camera assembly is mounted on a focusing stage (ASI LS-50). The filter wheel is mounted either between the objective and the tube lens (infinity space), or in front of the telecentric lens.

The excitation path is simplified compared to mesoSPIM v.5 but has the same effective excitation of NA=0.15. The left and right excitation arms are identical, each consisting of an optical fiber adapter, a collimator lens, an electrotuneable lens (*ETL*, Optotune EL-16-40-TC-VIS-5D-1-C), a 1:1 telescope (*L2, L3*, two Thorlabs AC254-100-A-ML), a galvo mirror (15 mm beam Thorlabs QS15X-AG), two folding mirrors, and an excitation objective (*L4*, modified Nikon 50 mm f/1.4 G objective), see **SI Fig. 14** and project github Wiki. In the excitation path, all 4 custom parts (folding mirror bracket, galvo mount, galvo heat sink, excitation objective mount, rendered in red) are machined from aluminium for thermal and mechanical stability. Switching of the laser between the arms is done at the laser combiner level (before the fiber coupling), with Oxxius L4Cc (405, 488, 561, 638 nm) laser combiner and with add-on module (product code “MDL-FSTM”) for fast switching between the two laser fibers. The galvo assembly is shown in **SI Video 12.**

As in its predecessor, the sample immersion chamber (**SI Fig. 15**) in the Benchtop mesoSPIM is stationary during image acquisition to ensure constant optical path for detection and excitation light. The immersion chamber (e.g., Portmann Instruments UG-753-H75 40×40×75 mm, fire-fused) is mounted on a kinematic table (Radiant Dyes RD-PDT-S) that allows tip, tilt and yaw adjustment of the chamber. The mounting is done with a custom-made chamber holder (3D printed from solvent-resistant plastic like PA12) and a magnetic kinematic base (Thorlabs KB1X1) that connects the holder with the kinematic table.

The sample is dipped into the immersion chamber and scanned through the light-sheet plane during the tile acquisition in a stepwise motion using the z-stage (ASI LS-50, 50 mm travel range with optical linear encoder and sub-micron accuracy). The vertical positioning between tiles is achieved with dual servo Y-stages (ASI LS-100 with 100 mm travel range), horizontal positioning with X-axis (ASI LS-50). Rotation of the sample is achieved with ASI C60-3060- SRS rotary stage.

The CAD design of mechanical components was performed in Autodesk Inventor 2023 software and the full model is available for download.

### Vibration absorption

Unlike previous mesoSPIM versions that were mounted on an air-floating optical table, the Benchtop mesoSPIM is built on a rigid but light breadboard (Thorlabs UltraLight Series II Breadboard, #PBG52522) resting on four Ø27.0 mm sorbothane isolator feet (Thorlabs, # AV4/M). The mass of the microscope on the breadboard is 25 kg (6.25 kg/isolator), so according to the isolator specifications, vibration frequencies higher than 20 Hz are well damped. To prevent low-frequency vibrations from people walking in the room, we recommend rooms with concrete floors and sturdy benches.

### Acquisition software

The user-friendly acquisition software^22^ allows full control via GUI and configuration files. It is written using the PyQt5 platform with multi-threading for high performance imaging (e.g., frame grabbing and file writing run in parallel threads), and it remains the same for all mesoSPIM versions. The Benchtop-specific code is limited to the configuration file, which is individual for each system. A high-level control software block diagram and GUI windows are shown in **SI Fig. 8**.

The electronics block diagram with the main computer-controlled components is shown in **SI Fig 16**. The waveforms generation is identical to the mesoSPIM v.5 (SI Figure 3 in Ref.^21^), but the number of National Instruments DAQmx boards was reduced to one (PXI-6733 with one BNC-2110 connector block) to minimize the footprint and cost. The system supports up to 4 laser lines, with both analog and digital modulation.

The software currently supports saving in 16-bit RAW (binary), Fiji^44^ TIFF, BigTIFF and Fiji BigDataViewer^45^ H5/XML file formats, along with necessary metadata files. For multi-tile/channel/illumination acquisitions the Fiji BigDataViewer H5/XML format is preferred because of fast writing speed and saving of rich metadata in XML, which makes the dataset ready for streamlined stitching and tiling in Fiji BigStitcher plugin^58^, independently of its size and RAM limitations.

### ASLM mode

The key feature of mesoSPIM which allows high and uniform z-resolution across large field of view is the axially swept light-sheet (ASLM) mode^21,23–25^. The ASLM is achieved in Benchtop the same way as in mesoSPIM v.5, by using an ETL (Optotune EL-16-40-TC-VIS-5D-1-C) in each excitation arm. The optimal ETL parameters (offset and amplitude) are easily adjusted by the user in the acquisition software, using parked beam (galvo scanning stopped) and laser scattering in the imaging medium, until a uniformly thin laser beam profile is achieved across the FOV in ASLM mode. This should be done in the absence of any sample in the FOV to avoid sample bleaching.

### Focus interpolation

For samples that are not well index matched with the imaging medium, or which do not use an immersion chamber, the focus distance depends on imaging plane z-position in the sample. Moving the sample in z changes its detection path optical distance and makes it go out of focus. This issue is tackled in the acquisition software, where the user can define two focus position values (beginning and end of the stack), and the software linearly interpolates between them during stack acquisition.

### Stitching, fusion, and 3D visualization

All datasets were saved in Fiji BigDataViewer H5/XML format, stitched in BigStitcher plugin^58^ and exported as either 16- or 8-bit TIFF files for visualization. Last column of **SI Table 3** shows the data size and number or tiles for each sample. Movies and screenshots were made in Imaris software (Oxford Instruments).

### Detection objective testing: contrast and field flatness

Our goal was to design a general test of detection objective properties which would be independent of the light-sheet illumination, to allow fast and reliable screening of detection objectives. In detection objective properties we are concerned about the overall contrast at high spatial frequencies, the uniformity of contrast across the field, and the field flatness.

A high-contrast square wave Ronchi ruling target glass slide (76.2 mm x 25.4 mm) with 40 line pairs per mm (lp/mm) pattern (Thorlabs R1L3S14N) was immersed in BK7 matching liquid (RI 1.5167, Cargille Cat #: 19586) inside a tall glass chamber. The total thickness of glass and BK7 matching liquid was adjusted to 20 mm to mimic the imaging conditions of a cleared specimen. The Ronchi ruling was illuminated from the back using a smartphone (Motorola g8) with a white screen image to provide uniform, diffuse, non-coherent illumination over a large FOV. The mesoSPIM camera exposure time was set to 50 ms to avoid temporal aliasing with the smartphone screen update rate.

During the acquisition, the chamber with Ronchi ruling was kept stationary, while the detection objective was moving in 10-µm steps, in a range from −300 to +300 µm from the best-focus position. Thus, a TIFF stack with 61 planes at various defocus positions was acquired. The analysis code splits each plane into a grid of subregions (32 x 20), and the contrast function is computed for each subregion:

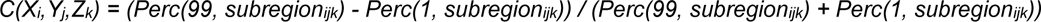

where *Perc* is the percentile function. Percentiles 1% and 99% were used instead of *min* and *max* respectively to account for noise.

The result is a 3D contrast map *C(x,y,z)* which is an estimate of modulation transfer function (MTF) at a given spatial frequency (40 lp/mm) for a given detection objective. The best-focus surface was computed from *C(x,y,z)* by taking its sections along *x* and *y* respectively, and measuring the sag (amplitude) of the high-intensity curve in each section (**SI Fig. 2c**, along *y-* axis). Note that *C(x,y,z)* is affected by the residual tip/tilt of the Ronchi test target, which is taken into account in the field flatness calculation.

By taking the maximum of *C(x,y,z)* along *z* we also computed the *maximum* contrast map *Cmax(x,y)* of the objective (**SI Fig. 2,3,4** left plot in each panel). This metric is independent from the tip/tilt of the Ronchi test target and can be used to cross-compare detection objectives.

The contrast map *C(x,y,z)* is also affected by the selection of tube lens, when infinity-corrected objectives are used. With Mitutoyo objectives we use standard MT-1 tube lens (f=200 mm), which is designed to provide a FOV (aka ‘image circle’ or ‘field number’) up to 30 mm when placed within nominal distance from the objective. With other objectives we used an Olympus wide-field tube lens SWTLU-C (f=180 mm, field number 26.5 mm). We noticed that mixing Mitutoyo objectives with SWTLU-C tube lens gave flatter field, but at a cost of 10% magnification reduction from nominal (by a factor 180/200 = 0.9).

### Chromatic effects on field flatness

To quantify the chromatic effects on detection objective contrast, we imaged the Ronchi slide (40 lp/mm) illuminated from behind (**SI Fig. 5b**) with a HAL100 tungsten-halogen lamp (Zeiss) fitted with a set of bandpass filters and a diffuser. The Ronchi slide was imaged in the air for control, and then in a chamber with DBE (17 mm DBE + 3 mm of glass between the slide and the detection objective). The slide was moved along Z axis at 10 µm steps between consecutive planes, thus emulating the conditions of a cleared tissue imaging.

The HAL100 lamp spectrum was balanced toward the blue range using a daylight blue filter (Olympus), which produced more uniform spread of intensity across the visible spectrum (**SI Fig. 5b**). Then, one of the following filters was applied to produce specific spectral bands: Chroma ET420/20, Semrock HQ470/30, Semrock Brightline HC 535/22, Semrock Brightline HC 595/31, Semrock Brightline Edge Basic 630/69 (combined with Edge Basic 594LP clean-up filter), Zeiss 697/75 (combined with Edge Basic 594LP). Lastly, the diffuser (120 grit, Thorlabs) was placed to ensure that light illuminates the Ronchi slide with a large angular spread. The illumination spectra were measured with the Hamamatsu VIS/NIR MiniSpectrometer model C10083CAH.

### Resolution measurement

Yellow-green fluorescent spheres (ex 441 nm, em 486 nm, Fluoresbrite® YG Microspheres 0.20 µm, Polysciences, Cat. # 17151) were mounted in 2% agarose in CUBIC-R+ medium (RI 1.52, TCI Chemicals, Prod. #T3741) inside a 10×10x45 mm glass cuvette. The sample cuvette was dipped in an immersion chamber filled with index-matched oil (RI 1.52, CUBIC mounting medium, TCI Chemicals, Prod. #M3294). Glass wall + oil thickness was 20 mm for Mitutoyo Plan Apo BD 5x/0.14 objective, and 5 mm for Mitutoyo 10x/0.28 and 20x/0.28 to control the spherical aberration effect but remain within realistic imaging conditions. The bead volume was acquired with 488-nm excitation laser and z-step = 1 µm.

From the resulting 3D stack of images, the bead centers were detected, filtered to exclude bead clusters, and bead image intensities were fitted with Gaussian function along x-y plane and z axis. The resulting PSF full width at half-maximum (FWHM) measurements were analysed visually by color-coding the the lateral resolution FWHM(x,y) and axial FWHM(z) across FOV, and by looking at their histogram, mean and standard deviation.

As seen in **SI Fig 1** histograms, the axial resolution of Benchtop is 3.3-4.0 µm at all magnifications (determined by the effective light-sheet thickness in ASLM mode, which is independent of the detection objective), and the lateral resolution ranges from 1.5 µm (Mitutoyo G Plan Apo 20x/0.28-t3.5) to 2.7 µm (Mitutoyo BD Plan Apo 5x/0.14), depending on the objective NA and the medium thickness used in the test. **Sample mounting** iDISCO and ECi cleared samples were either placed in a dipping cuvette or clamped with adjustable 3D printed clamps (**Fig. 1f**) and dipped into an immersion chamber (**SI Fig. 14**) containing DBE or ECi, respectively. Whole mouse body cleared with vDISCO (**Fig. 2a**) was placed into BABB-filled rectangular cuvette (Portmann Instruments UG-751-H80, 25×25x80mm) which was held on top by metal clamp (Thorlabs BSH1/M) and imaged directly without dipping into immersion chamber, with focus interpolation.

All our sample holders were 3D printed from PA12 (polyamide) using industrial SLS printer at the UZH AMF BIOC facility.

### Video tutorials

https://www.youtube.com/@mesoSPIM

### Viral injection and tissue collection

All mouse experiments were conducted in accordance with standard ethical guidelines and the guidelines from the Veterinary Office of Switzerland and were approved by the Zurich Cantonal Veterinary Office.

For sparse retrograde viral labelling, we stereotactically injected an eight months old male Vglut2-Cre mouse. The mouse was briefly anesthetized with isoflurane (2%) in oxygen in an anesthesia chamber and subsequently immobilized for intracerebral injection in a stereotactic frame (Kopf Instruments). Body temperature was constantly maintained at ∼37°C. To prevent drying, the mouse eyes were covered with vitamin A containing cream (Bausch & Lomb). Anesthesia was maintained at 1% isoflurane in oxygen. We stereotactically injected ∼200nl of a retro-grade AAV encoding a cre-dependent variant of td-Tomato(ssAAV-retro/2-CAG-dlox-tdTomato(rev)-dlox-WPRE-bGHp(A); concentration: 0.85 x 10E12 vg/ml; UZH Viral Vector Facility) in the right LHb using the following coordinates from bregma (in mm): −1.7 anteroposterior (AP), +0.5 mediolateral (ML), −2.3 dorsoventral (DV).

### iDISCO mouse brain clearing

Mouse brains were stained for different neuronal cell types or amyloid plaques and arterial vessels and cleared with a modified version of the iDISCO protocol^47^. For AAV-injected Vglut2-Cre mice, animals were used after 28 days of expression. For Thy1-positive neuronal labeling, B6.Cg-Tg(Thy1-YFP)16Jrs/J (#003709) mice were used. For plaque and arterial vessel labeling, B6.Cg-Tg(Thy1-APPSw,Thy1-PSEN1*L166P)21Jckr were used. All mice were deeply anesthetized with ketamine (100 mg/kg body weight; Streuli Pharma AG) and transcardially perfused with ice-cold phosphate buffer, pH 7.4 (PBS), followed by ice-cold 4% paraformaldehyde (PFA) in PBS. After 4.5 hours of post-fixation at 4°C shaking at 40 rpm, brains were washed with PBS for three times at room temperature (RT), with daily solution exchange.

Samples were dehydrated in serial incubations of 20%, 40%, 60%, 80% methanol (MeOH) in ddH2O, followed by 2 times 100% MeOH, each for 1 hour at RT and 40 rpm. Pre-clearing was performed in 33% MeOH in dichloromethane (DCM) overnight (o.n.) at RT and 40 rpm. After 2 times washing in 100% MeOH each for 1 hour at RT and then 4°C at 40 rpm, bleaching was performed in 5% hydrogen peroxide in MeOH for 20 hours at 4°C and 40 rpm. Samples were rehydrated in serial incubations of 80%, 60%, 40%, and 20% MeOH in in ddH2O, followed by PBS, each for 1 hour at RT and 40 rpm. Permeabilization was performed by incubating the mouse brains 2 times in 0.2% TritonX-100 in PBS each for 1 hour at RT and 40 rpm, followed by incubation in 0.2% TritonX-100 + 10% dimethyl sulfoxide (DMSO) + 2.3% glycine + 0.1% sodium azide (NaN3) in PBS for 3 days at 37°C and 65 rpm. Blocking was performed in 0.2% Tween-20 + 0.1% heparine (10 mg/ml) + 5% DMSO + 6% donkey serum in PBS for 2 days at 37°C and 65 rpm. All samples were stained gradually with the respective primary and secondary antibodies listed below in 0.2% Tween-20 + 0.1% heparine + 5% DMSO + 0.1% NaN3 in PBS (staining buffer) in a total volume of 1.5 ml per sample every week for 4 weeks at 37°C and 65 rpm. Between first and secondary antibody incubation, samples were washed in staining buffer 4 times each for one hour followed by 5- th time o.n. at RT.

The following antibodies were used. For Vglut2-Cre mice, the primary polyclonal rabbit-anti-RFP antibody (Rockland, 600-401-379-RTU, dilution 1:2000) and secondary donkey-anti-rabbit-Cy3 antibody (Jackson ImmunoResearch, 711-165-152, dilution 1:2000) were used. For Thy1-YFP mice, the primary polyclonal chicken-anti-GFP antibody (Aves Labs, GFP- 1020, dilution 1:400) and secondary donkey-anti-chicken-AlexaFluor594 antibody (Jackson Immuno Research, 703-585-155, dilution 1:400) were used. For APP/PS1 mice, the dye hFTAA^59^ was used at dilution of 1:400 in combination with the Cy3-conjugated monoclonal anti-α-smooth muscle actin antibody (Sigma Aldrich, C6198) at dilution of 1:800.

After staining, mouse brains were washed in staining buffer 4 times each for one hour and 2 times o.n. at RT. Samples were dehydrated in serial incubations of 20%, 40%, 60%, 80% MeOH in ddH2O, followed by 2 times 100% MeOH, each for 1 hour at RT and 40 rpm. Clearing was performed in 33% MeOH in DCM o.n. at RT and 40 rpm, followed by incubation in 100% DCM 2 times each for 30 minutes. Refractive index matching (RI = 1.56) was achieved in dibenzylether (DBE).

### vDISCO whole mouse body clearing

Animal experiments of this part followed European directive 2010/63/EU for animal research, reported according to the Animal Research: Reporting of In Vivo Experiments (ARRIVE) criteria, complied with the ‘3Rs’ measure and were approved by the ethical review board of the government of Upper Bavaria (Regierung von Oberbayern, Munich, Germany) and conformed to institutional guidelines of Klinikum der Universität München/Ludwig Maximilian University of Munich). The severity of the procedure was low.

The Thy1-GFP line M (2.5 month old) mouse, from which the brain and the spinal cord were dissected out, and the CX3CR1-GFP (P14) mouse were stained and cleared with vDISCO^60^. The detailed vDISCO protocol is available in Ref.^27^ Briefly, the 0.01M PBS (1x PBS) perfused and 4% paraformaldehyde (Morphisto, 11762.01000) fixed mouse body was placed in a 300 ml glass chamber (Omnilab, 5163279) filled with 250–300 ml of solution, which covered the body completely. Next, a perfusion setting able to pump the solution in the glass chamber into the transcardial circulatory system of the mouse in recycling manner was established involving a peristaltic pump (ISMATEC, REGLO Digital MS-4/8 ISM 834; reference tubing, SC0266) and two tubing channels: the first was set to circulate the solution through the heart into the vasculature by using a perfusion needle (Leica, 39471024), the second was immersed into the solution chamber where the animal was placed. To fix the needle tip in place and to ensure extensive perfusion, a drop of superglue (Pattex, PSK1C) was added onto the hole of the heart where the needle was inserted. By using this perfusion setting, the animal was first perfused with a decolorization solution for 2 days at room temperature, refreshing the solution when turned yellow, and then by a decalcification solution for 2 days at room temperature. In between these two solutions, a shorter step that consisted of perfusing the body with 1x PBS for 3 hours 3 times at room temperature was performed as washing step. The decolorization solution was made with 25–30 vol% dilution of CUBIC reagent #1 (Ref.^61^): 25 wt% urea (Carl Roth, 3941.3), 25 wt% N,N,N′,N′-tetrakis (2- hydroxypropyl)ethylenediamine (Sigma-Aldrich, 122262) and 15 wt% Triton X-100 (AppliChem, A4975,1000) in 1x PBS. The decalcification solution consisted of 10 wt/vol% EDTA (Carl Roth, 1702922685) in 1x PBS, pH 8–9 adjusted with sodium hydroxide (Sigma-Aldrich, 71687).

Next, the mouse was perfused with 250 ml of permeabilization solution consisting of 1.5% goat serum (Gibco, 16210072), 0.5% Triton X-100, 0.5 mM of methyl-β-cyclodextrin (Sigma, 332615), 0.2% trans-1-acetyl-4-hydroxy-l-proline (Sigma-Aldrich, 441562) and 0.05% sodium azide (Sigma-Aldrich, 71290) in 1x PBS for half a day at room temperature. After this, the mouse was perfused for 6 days with 250 ml of the same permeabilization solution containing 290 μl of propidium iodide (PI, stock concentration 1 mg ml^−1^ Sigma-Aldrich, P4864) and 35 μl of Atto647N conjugated anti-GFP nanobooster (Chromotek, gba647n- 100). When the staining step was completed, the mouse was washed by perfusing with the washing solution (1.5% goat serum, 0.5% Triton X-100, 0.05% of sodium azide in 1x PBS) for 3 hours 3 times at room temperature and with 1x PBS for 3 hours 3 times at room temperature. Finally, the mouse was cleared in the glass chamber as follows: with passive incubation at room temperature in a gradient of tetrahydrofuran (Sigma-Aldrich, 186562)/double-distilled water (50, 70, 80, 100 and again 100 vol%) 12 hours each step, followed by 3 hours in dichloromethane (Sigma-Aldrich, 270997) and in the end in a mixture of benzyl alcohol (Sigma-Aldrich, 24122) and benzyl benzoate (Sigma-Aldrich, W213802) 1:2. During all incubation steps, the glass chamber was sealed with parafilm and covered with aluminum foil. All the clearing steps were performed in a fume hood.

### Human brain tissue preparation

The body donors were provided by the body donation program of the Department of Anatomy and Embryology, Maastricht University. The tissue donor gave their informed and written consent to the donation of their body for teaching and research purposes as regulated by the Dutch law for the use of human remains for scientific research and education (“Wet op de Lijkbezorging”). Accordingly, a handwritten and signed codicil from the donor posed when still alive and well, is kept at the Department of Anatomy and Embryology Faculty of Health, Medicine and Life Sciences, Maastricht University, Maastricht, The Netherlands. Three human occipital lobe samples were obtained from 2 body donors each (Occipital lobe 1: 101-year-old female; Occipital lobe 2: 90 year old male; no known neurological disease, respectively). All six samples were taken approx. 3 cm anterior to the occipital pole and around the V1/V2 border (see **SI Fig. 9 and 10**). Brains were first fixed in situ by full body perfusion via the femoral artery. Under a pressure of 0.2 bar the body was perfused by 10 l fixation fluid (1.8 vol% formaldehyde, 20% ethanol, 8.4% glycerine in water) within 1.5–2 hours. Thereafter, the body was preserved at least 4 weeks for post-fixation submersed in the same fluid. Subsequently, brains were recovered by calvarial dissection and stored in 4% paraformaldehyde in 0.1 M phosphate buffered saline (PBS) until further processing.

The samples from Occipital lobe 1 were stained with MASH-MB and Occipital lobe 2 samples with MASH-NR, as previously described^48^, with minor adjustments to the original protocol: The tissue was dehydrated in 20, 40, 60, 80, 100% methanol (MeOH) in distilled water for 1h each at room temperature (RT), followed by 1h in 100% MeOH and overnight bleaching in 5% H2O2 in MeOH at 4°C. Samples were then rehydrated in 80, 60, 40, 20% MeOH, permeabilized 2x for 1h in PBS containing 0.2% Triton X-100 (PBST). This was followed by a second bleaching step in freshly filtered 50% aqueous potassium disulfite solution. The samples were then thoroughly rinsed in distilled water 5x and washed for another 1h. For staining, the samples were incubated in 0.001% methylene blue (Occipital lobe 1) or 0.001% neutral red (Occipital lobe 2) solution in McIlvain buffer (phosphate-citrate buffer)^62^ at pH 4 for 5 days at room temperature. After 2.5 days the samples were flipped to allow for equal penetration of the dye from both sides. After staining, samples were washed 2x for 1h in the same buffer solution, dehydrated in 20, 40, 60, 80, 2x 100% MeOH for 1h each, and delipidated in 66% dichloromethane (DCM)/33% MeOH overnight. This was followed by 2x 1h washes in 100% DCM and immersion in ethyl cinnamate (ECi)^63^.

### *Xenopus* tropicalis sample preparation

All tadpole experiments were conducted in accordance with standard ethical guidelines and the guidelines from the Veterinary Office of Switzerland and were approved by the Zurich Cantonal Veterinary Office.

Whole-mount *Xenopus* immunofluorescence was adapted from a previously described^17^. A stage 58 *Xenopus tropicalis* tadpole was fixed overnight in 4% PFA and dehydrated with 100% MeOH for 3 times 24h. Bleaching was performed under room lightning conditions in 10% H2O2 / 23% H2O / 66% MeOH for 8 days, replacing the bleaching solution every two days until fully bleached.

The tadpole was stepwise rehydrated in 1x PBS with 0.1% Triton X-100 (PBST) and blocking was performed for 3 days at 4°C in 10% CAS-Block / 90% PBST (008120, Life Technologies). Staining was performed using Atp1a1 antibody (1:500, DSHB, a5) diluted in 100% CAS-Block for 5 days at 4°C on a rotating wheel. The tadpole was washed 3 times for 8h and 2 times for 1 day with PBST at RT, blocked again for 1 day at 4°C (10% CAS-Block / 90% PBST) and incubated 72h at 4°C on a rotating wheel with the secondary antibody (1:500, Alexa-Fluor-555, 405324 P4U/BioLegend UK Ltd) diluted in 100% CAS-Block. The tadpole was washed for 3x 8h and 2x 1 day with PBST.

The tadpole was dehydrated in 25% MeOH/75% 1x PBS (1 day), 50% MeOH/50% 1x PBS (1 day), 75% MeOH/25% 1x PBS (1 day), three times 100% MeOH (2 times 8h, 1 time 24h). Finally, clearing was performed in BABB (benzyl alcohol:benzyl benzoate 1:2, (Merck 8187011000 and Sigma 108006) for 48h.

For samples imaged on the SPIM-tower, BMS-453-treated (1 μM final concentration, 0.1%DMSO) and untreated embryos (0.1% DMSO) were fixed at respectively st. 33, 37 and 42, stained and cleared as described before^17^. In brief, the primary antibody mix was anti-Atp1a1 (1:200, DSHB, A5), anti-col2a1 (1:200, DSHB, II-II6B3) and anti-TNNT2 (1:200, DSHB, CT3). The secondary antibody was Alexa Fluor 555 goat anti-mouse IgG (minimal x-reactivity) (405324, P4U/BioLegend UK). For nuclear counterstaining, DAPI (20 μg/ml, ThermoFisher, D1306) was added to the primary antibody mixture.

### Chicken embryo

All chicken embryo experiments were conducted in accordance with standard ethical guidelines and the guidelines from the Veterinary Office of Switzerland and were approved by the Zurich Cantonal Veterinary Office.

Neurofilament staining of a whole-mount chicken embryo: The embryo was sacrificed at day 9 of development and fixed in 4% paraformaldehyde for 3.5 hours at room temperature. For best results, the embryo was kept in constant, gentle motion throughout the staining procedure. Incubations were at 4°C if not declared differently. The tissue was permeabilized in 1% Triton X-100/PBS for 30 hours, followed by an incubation in 20 mM lysine in 0.1 M sodium phosphate, pH 7.3 for 18 hours. The embryo was rinsed with five changes of PBS containing 0.2% Triton. To avoid unspecific antibody binding the embryo was incubated in 10% FCS (fetal calf serum) in PBS for 48 hours. The primary antibody mouse anti neurofilament (1:1’500, RMO270, Invitrogen 13-0700) was added for 60 hours. Unbound primary antibody was removed by ten changes of 0.2% Triton/PBS and an additional incubation overnight. After re-blocking in 10% FCS/PBS for 48 hours, the embryo was incubated with the secondary antibody goat anti-mouse IgG-Cy3 (1:1’000, Jackson ImmunoResearch 115-165-003) for 48 hours. Afterwards, the embryo was washed ten times with 0.2% Triton/PBS followed by incubation overnight. For imaging, the tissue was dehydrated in a methanol gradient (25%, 50%, 75% in H2O and 2x 100%, minimum 4 hours each step at room temperature) and cleared using 1:2 benzyl alcohol: benzyl benzoate (BABB) solution (again gentle shaking is recommended for dehydration and clearing). The tissue and staining are stable for months when kept at 4°C in the dark.

### CaF2 crystal preparation

Four 1cm^3^ CaF2 crystals, all sides polished, were acquired from Crystran. The shape and polish of the crystals were chosen to minimize multiple scattering of light in the sample. Two of these crystals were irradiated in a ^60^Co irradiator at Penn State University. The crystals received dosages of ∼100 kRad and 5 MRad each. As the radiation source was isotropic and the attenuation of 1.17 MeV and 1.33 MeV gamma-rays by 1 centimeter of CaF2 is low^64^, the deposition of energy by the gamma-rays in the crystal is expected to be uniform. The two other crystals were left without irradiation for comparison (*blank* references).

The crystals were placed in a custom-made sample holder (**Sl Fig. 12b**) and were imaged without immersion both before and after gamma-ray irradiation. By comparing the images before and after irradiation (as well as irradiated vs *blank*), we demonstrate that the mesoSPIM can clearly measure fluorescence from color centers induced by irradiation. **Fig. 3i** shows the images in response to the 405 nm laser. The response at different laser settings, comparison to blank, and distribution of color centers are shown in **SI Fig. 12** and discussed in **Sl Note 2**.

## Data availability

Representative microscopy data are available in the BioStudies database (https://www.ebi.ac.uk/biostudies/) under accession number S-BIAD963.

## Code availability

The Benchtop design files, list of parts, and building instructions (Wiki): https://github.com/mesoSPIM/benchtop-hardware The mesoSPIM control software: https://github.com/mesoSPIM/mesoSPIM-control The detection objective testing, including chromatic effects on field flatness: https://github.com/nvladimus/lens-testing The PSF quantification (ETL-on and-off cases): https://github.com/mesoSPIM/mesoSPIM-PSFanalysis The optical simulations in Optalix software: https://github.com/mesoSPIM/optical-simulations

## Acknowledgments

The authors thank Stefan Giger from the Brain Research Institute for mechanical prototyping, Fabian Eggiman, Lukas Lüchinger and Sascha Weidner from the Additive Manufacturing Facility (AMF) of the University of Zurich (UZH)for help with designing and 3D printing of custom sample holders, Daniel Invernot, Melanie Horn and Julia Traversari from UZH for *Xenopus* animal care and husbandry, Andrew Woehler from HHMI Janelia JET for providing Thorlabs super apochromat objectives, Giulia Miracca from the University Hospital Zurich for requesting and testing BSL-3 cuvette clamps; Igor Jovanovic from the University of Michigan for the irradiation of crystals, Michel Rickhaus and Joseph Woods from UZH for the use of the fluorescence spectrophotometer, and Celine Heeb and Lorenz Weber for help with Wiki instructions on building a Benchtop mesoSPIM. Finally, we thank our anonymous reviewers for asking good questions that helped improve the manuscript.

## Supplementary Videos

**SI Video 1** Contrast map of Olympus XLFluor 4x/0.28 objective based on imaging a 40lp/mm Ronchi grid through 20 mm of high-index medium (RI 1.52). The Z axis represents focus depth F [µm] scaled by 10x for demonstration purpose.

**SI Video 2** Mouse brain with spinal cord dissected from a 2.5-month-old Thy1-GFP line M animal, cleared with vDISCO and imaged at 5x magnification.

**SI Video 3** Whole mouse P14 stained with propidium iodide and cleared with vDISCO, imaged at 0.9x magnification.

**SI Video 4** Chicken embryo at E9 stage stained with M Anti-RMO270, G Anti-Mouse Cy3, cleared with BABB, imaged at 1.2x magnification.

**SI Video 5** Brain of APPPS-1 mouse, stained with hFTAA (amyloid plaques) and SMA-Cy3 (arterial vessels), cleared with iDISCO, imaged at 1.2x magnification.

**SI Video 6** Brain of Vglut2-Cre mouse with retrograde AAV cre-dependent variant of td-Tomato (ssAAV-retro/2-CAG-dlox-tdTomato (rev)-dlox-WPRE-bGHp(A), cleared with iDISCO, imaged at 4x magnification.

**SI Video 7** One hemisphere of Thy1-YFP AF-561 mouse, cleared with iDISCO, imaged at 5x magnification.

**SI Video 8** Piece of human cortex (occipital lobe V2) from 90 y.o. male donor (Occipital lobe 2), stained with neutral red and cleared with the MASH-NR protocol (Nissl staining with neutral red; ECi as immersion medium), imaged at 5x magnification.

**SI Video 9** Human cortex (part of occipital lobe V2) from 101 y.o. female donor (Occipital lobe 1), stained with methylene blue and cleared with the MASH-NR MB protocol (Nissl staining with methylene blue; ECi as immersion medium), imaged at 5x magnification.

**SI Video 10** Xenopus tadpole stage 58, stained with Atp1a1 AF-555, cleared with BABB, imaged at 5x.

**SI Video 11** Benchtop mesoSPIM assembly from modules in 50 minutes at EMBO Course “3D developmental imaging”, Oeiras, 2022. URL: https://youtu.be/4nsc5BLfjYc

**SI Video 12** Galvo assembly for Benchtop mesoSPIM excitation arm. URL: https://youtu.be/53Eq5iCVMGg

## Supplementary notes

### SI Note 1: Acquisition speed and file size

To image 1 cm^3^ (dimensions *Sx = Sy = Sz* = 10 mm) of sample in multi-tiling acquisition, the number of frames along each dimension are

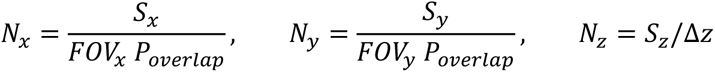

where *FOVx* and *FOVy* are field of view dimensions on the sample side (camera chip dimensions divided by the system magnification), *Poverlap* = 0.9 is the tile overlap ratio in X and Y used for stitching (10%, default setting), Δ*z* is the z-step between planes.

The total acquisition time is

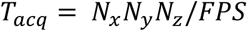

where FPS is the effective frame rate (frames/s).

Fast mode. In this mode one can choose under-sampling in Z while still maintaining sufficient resolution for axons and dendrites detection.

Nyquist mode. This mode samples the axial PSF of the system at the optimal step, about 2 planes per PSF axial size, FWHMz.

The code for calculating the acquisition speed and file size is available as Python notebook: https://github.com/mesoSPIM/image-processing/blob/main/notebooks/volumetric-speed-calculation.ipynb

### SI Note 2: Imaging of color centers for particle detectors

In the main text, we present a Benchtop mesoSPIM application for imaging image color centers induced in crystals by irradiation. This application has the ultimate goal of dark matter and neutrino detection, which are relevant topics in the fields of particle physics, nuclear engineering and nuclear non-proliferation safeguards.

The objective of this section is to provide the reader outside of these fields with: 1) the background on the search and detection of elusive particles, such as dark matter and neutrinos, as well as the application of neutrino detectors as nuclear non-proliferation safeguards, 2) additional information on the topic of microscopy imaging of color centers (and its use for particle detectors), 3) andoverview the current research and ongoing developments, as well as future prospects.

### Color centers signals from particle interactions

#### Dark matter

There is clear evidence for the existence of dark matter^1,2^ -- a type of matter that exerts a gravitational pull in our galaxy (and universe), but is ‘dark’ because it does not interact with photons in the same way ordinary matter does. Although five times more abundant than ordinary matter^1^, dark matter particles have not yet been detected. Many theories predict it to be a weakly interacting massive particle. As our solar system is immersed in a dark matter sea and travels across it at 220 km/s, we should constantly receive a large flux from these particles. Although most of these particles should travel across the Earth unnoticed, we expect them to interact occasionally with the nuclei of ordinary atoms, making them recoil. Many experiments have been searching for signals from these dark matter collisions with nuclei^3–9^. Most of them rely on the collection of charge, prompt scintillation photons or phonons as dark-matter induced signals. In this work, we present the first tests done with the Benchtop mesoSPIM using a new approach, proposed by Cogswell et al.^10^: the use of SPIM to detect color centers signals, induced by the collision of particles, such as dark matter, with atoms in the lattice of a crystal.

A schematic of how dark matter (and other particles) could create color centers in a crystal is shown in SI Notes Fig. 1. For a detailed review of the concept, we refer the reader to Refs^10–12^. For a detailed description of color centers, we refer the reader to Ref.^13^

#### Nuclear reactor neutrinos

To detect another elusive particle, the neutrino, ton-sized detectors or powerful accelerators are usually required to obtain a few events per day^15–17^. While elastic collisions of neutrinos with nuclei have just recently been observed using accelerator neutrino sources^16^, these interactions from reactor neutrinos have not yet been detected. The challenge is to bring a very sensitive detector close enough to the reactor. Small passive detectors, such as transparent crystals, could be more easily placed close to the reactor and work as nuclear-nonproliferation safeguards^10^. In this application, crystals would be exposed to the high flux of neutrinos from the reactor and scanned with SPIM (*ex-situ*) to image neutrino-induced color centers. By counting the number of new centers in the crystal, it would be possible to understand the reactor power^10^. This knowledge could be then used to estimate (or exclude) the production of plutonium in the monitored reactor^10^.

#### Imaging of color centers

While the interaction of elusive particles (W, *v*) with the crystal lattice cause atom dislocations of only a few nanometers (SI Notes Fig. 1), color centers can be probed with fluorescence microscopy at optical wavelengths. The fluorescence response of color centers enables faster and less expensive imaging than other techniques used to probe nm-sized features, such as transmission electron microscopy (TEM) and atomic force microscopy (AFM).

The imaging of single color centers has already been performed with widefield and confocal fluorescence microscopes^18,19^. Imaging these color centers with SPIM offers further advantages: good optical sectioning compared to widefield imaging, less bleaching, higher throughput, and larger maximum sample size compared to confocal imaging.

The non-destructive, fast and isotropic scan of large 3D-volumes offered by mesoSPIM is thus well suited for imaging large amounts of color-center based particle detectors. Another possible application of color center imaging with the mesoSPIM is fission track dating of transparent rocks: the mesoSPIM could offer a fast imaging method for large amounts of minerals without the need of etching, as described in Ref^12^.

#### Benchtop mesoSPIM imaging of particle-induced color centers in CaF2

In this work, we show the first tests of imaging particle-induced color centers with the mesoSPIM. This is part of the international collaborative work of PALEOCCENE (Passive low-energy optical color center nuclear recoil) – a group of scientists working on the R&D of passive detectors of dark matter and neutrinos at low energy thresholds^11^.

For the purpose of initial tests, we acquired transparent CaF2 crystals, irradiated them, and imaged them with the mesoSPIM, as described in the Methods section. The Sl Fig 10a shows the crystal ready for imaging before and while the light sheet is on. A custom-made crystal holder, shown in b was made for keeping the samples in a stable position and allowing for changing crystals in consecutive scans without the need of complete refocusing. The fluorescence of an irradiated crystal in response to the 405 nm light sheet is clearly distinguished from the background, as shown by the contrast of the illuminated area in c and the pixel intensity distribution in c. While the fluorescence is mostly homogeneous across the crystal (presumably from many nm-sized color centers), a few clusters of high-intensity pixels appear at small scales (∼10 µm). These structures are especially clear when imaged with the 20x/0.28 objective. A few examples are shown in d.

The structures shown in Sl Fig.10d are identified by an algorithm, which scans the area of the illuminated ROI and outputs the contours of clusters of high-intensity pixels. The parameters used to define the selection thresholds of intensity and clustering are preliminary and will be benchmarked and improved in the future with data from dedicated ion-irradiation (track-inducing) campaigns.

To verify whether the selected structures are intrinsic to the crystal – and not some random noise from the camera – repeated scans are compared. The matching correlation between them is calculated with the skimage.feature.template_matching method, where the smallest rectangle enclosing the feature with N pixels is compared to the repeated scan using a normalized cross-correlation. The result ranges from −1 to 1, where 1 represents a perfect match. Most of the values are around zero, which is expected from random matching of any given group of N pixels which do not contain structures. The first two examples of the identified structures shown in Sl Fig.10d are observed at the same respective spots in the repeated scans and thus display large matching correlation coefficients (shown in red in e). Their matching coefficients are 7.9 and 7.3 sigma away from the mean of their respective distributions. The other values similar to the ones in red are due to the matching with neighboring z-planes, as the imaged structures span across a few z-scans. These distributions vary as the N number of pixels in the rectangles also vary.

While this analysis shows that these “track-like” structures are intrinsic fluorescent features in the crystal, the origin of these features is not yet clear. One possibility is that the passage of particles such as cosmic rays may have created a track of color centers. Cosmic rays hit the Earth constantly, especially at high altitudes. The crystals were likely exposed to this natural irradiation since their production and more intensively while they were mailed by air after the purchase and for the irradiation campaigns. Despite the unknown origin of these features, this study demonstrates the capability of the benchtop mesoSPIM in identifying “track-like” structures of color centers. While the tracks from dark matter or neutrino interactions would have sizes below the resolution power of the mesoSPIM, understanding track formation is part of the R&D process of the concept. Furthermore, “the energy reconstruction of dark matter / neutrino interactions may be possible through the larger intensity of fluorescence (pixel brightness) measured from a region containing a full track in comparison to a single-site color center”^12^. The improved resolution provided at 20x magnification will be especially relevant when imaging single color centers or single tracks from nuclear recoils produced by neutrons – a proof-of-concept test planned as a next step.

With the mesoSPIM images of the crystals irradiated with gamma rays, we performed further analyses with the aim of understanding: i) the distribution/homogeneity of color centers at larger scales (across milliliters of material), ii) the sources of background, and iii) the color of the imaged color centers, and how the results obtained with the mesoSPIM compare to far-field spectroscopy.

To quantify the level and homogeneity of fluorescence from blank and irradiated crystals, we imaged them at 1x magnification and calculated the average pixel intensity from every z-image of 350 scans taken at 10 µm steps inside the crystals. The obtained mean values and the respective gaussian standard deviations are shown in Sl Fig 10f. For this estimation, only pixels inside the illumination ROI #1 and at a distance larger than ∼0.2 mm from the surface were selected to avoid surface background (as we found that the surface of blank crystals present a slightly higher signal intensity than their bulk). Possible sources of the higher surface fluorescents include dust or any other residual material which is auto-fluorescent and difficult to remove by usual cleaning methods, such as machine oils or surface coating, depending on the manufacturing of the crystal. The bulk of *blank* (not irradiated or before irradiation) crystals displayed low net fluorescence level, as observed in Sl Fig.10f: the fluorescence intensities in response to all the selected laser excitations were very close to the background level, which was estimated with the laser off.

By imaging these crystals in the mesoSPIM with different laser excitations (405, 488, and 561 nm) and filters (quadrupole or long pass), we can understand the colors / wavelengths absorbed and reemitted as well as verify whether the signal corresponds to fluorescence or Raman scattering. Sl Fig.10f shows that irradiated crystals absorb 405 nm light and fluoresce in the blue: the most intense emission is in response to 405 nm light and faints when the blue spectrum is cut by a 515 nm longpass filter. This blue fluorescence has been confirmed by the absorption and emission spectra measured in response to light from 250 to 800 nm with an Edinburgh Instruments FS5 spectrofluorometer. An example of the measured fluorescence from an irradiated crystal in response to 400 ± 10 nm light is shown in Sl Fig.10g. The response to shorter excitation wavelengths (down to ∼340 nm) consistently presents the emission peak at ∼ 425 nm. This resuult validates the observations obtained with the mesoSPIM: the irradiated crystals fluoresce in blue and the signal is not Raman scattering. A fainter emission peak at ∼740 nm is also observed and its intensity increases at larger excitation wavelengths (∼600 nm). This also matches the results obtained with the mesoSPIM: a significant fluorescent signal in response to 561-nm excitation light.

While the mesoSPIM data agrees with the far-field spectroscopy, there are several advantages offered by the mesoSPIM. With the sectioned images, we can define an ROI avoiding surface background, estimate the fluorescence colors as well as quantify the distribution of color centers across the sample. The mesoSPIM can also image single clusters of color centers and precisely identify their position, a feature that is crucial for the concept of particle detectors discussed in this section.

### SI Note 3: Effects of spherical aberration on excitation beam

The presence of a thick (10-30 mm) layer of immersion medium is expected to broaden the excitation beam and the detection PSF due to spherical aberration. To estimate these effects, we simulated a simplified system in OptalixPro software (**SI Notes Fig. 2**), with excitation objective simulated as ideal lens (f=50 mm) and with chamber glass walls neglected for simplicity (their refractive index 1.52 is close to the imaging medium 1.56). From the **SI Table 9** one can see that the beam diameter slowly increases with medium thickness up to 20 mm, then transiently decreases at thickness 30 mm, and then increases again at thickness 40 mm. In practical mesoSPIM usage, the beam is expected to remain in the range between 2.4 and 3.0 µm (medium thickness 10-30 mm).

In the absence of ASLM scanning (ETL amplitude set to 0) we experimentally measured the beam waist diameter FWHM to be 2.8±0.33 µm (mean±std, n=103 beads) for the 488 nm excitation, which agrees almost perfectly with the simulations. The experimental results are available at notebook https://github.com/mesoSPIM/mesoSPIM-PSFanalysis/benchtop-MitutoyoBD-5X(TL-MT1)-ETL-off.ipynb.

Note that in the ASLM-on mode the beam is scanned axially, so that the beam experiences a range of medium thicknesses during scanning, which effectively makes it broader, in agreement with our experimental measurements of beads in ASLM regime, which show field-averaged PSF FWHM(z) in the range of 3.3-4.0 µm (SI Fig.1), a spread factor of about 37% compared to the ASLM-off simulated and experimentally measured beam diameter.

These simulations suggest that Nikon 50 mm f/1.4 G excitation objectives achieve close to diffraction limited performance for NA=0.15 beam focusing, and that broadening of the excitation beam is mostly due to the immersion medium.

### SI Note 4: Effects of spherical aberration on detection performance

To evaluate the effect of spherical aberration (caused by the immersion medium) on the *detection* performance we simulated a simplified system: a 20-mm thick medium with a given R.I., an ideal 5x objective (f=40 mm) and an ideal tube lens (f=200 mm). The performance was measured by PSF properties such as FWHM(x,y) and Strehl ratio. While FWHM is often used to estimate the resolution, Strehl ratio estimates how much PSF intensity peak is *reduced* compared to an ideal aberration-free PSF, which affects the image contrast. An optical system with a Strehl ratio above 0.8 is considered diffraction-limited.

In agreement with our experimental results (SI Fig. 2), the NA of the simulated detection objective has a strong impact on the imaging performance, especially on the contrast. With NA changing from 0.15 to 0.30, the PSF FWHM(x,y) remained nearly constant (thus cancelling the positive effect of higher NA), while the Strehl ratio dropped from 0.65 to 0.04 (SI Table 10), which negatively affects image contrast.

At a fixed medium thickness (20 mm) and objective NA (0.15), variation of medium refractive index in a range from 1.47 to 1.99 did not result any significant deterioration of either PSF size or Strehl ratio (SI Table 11).

At a fixed medium R.I. and objective NA, the medium thickness itself had a strong impact on system’s performance, such that Strehl ratio was nearly diffraction-limited at medium thickness 5-15 mm and deteriorated to 0.40 at thickness 30 mm (SI Table 12), in agreement with our experience that image contrast drops when the sample and medium are thick, even for well cleared samples. We therefore remind the users that sample and medium thickness should be kept as small as possible for optimal results.

---

**SI Fig. 1.**
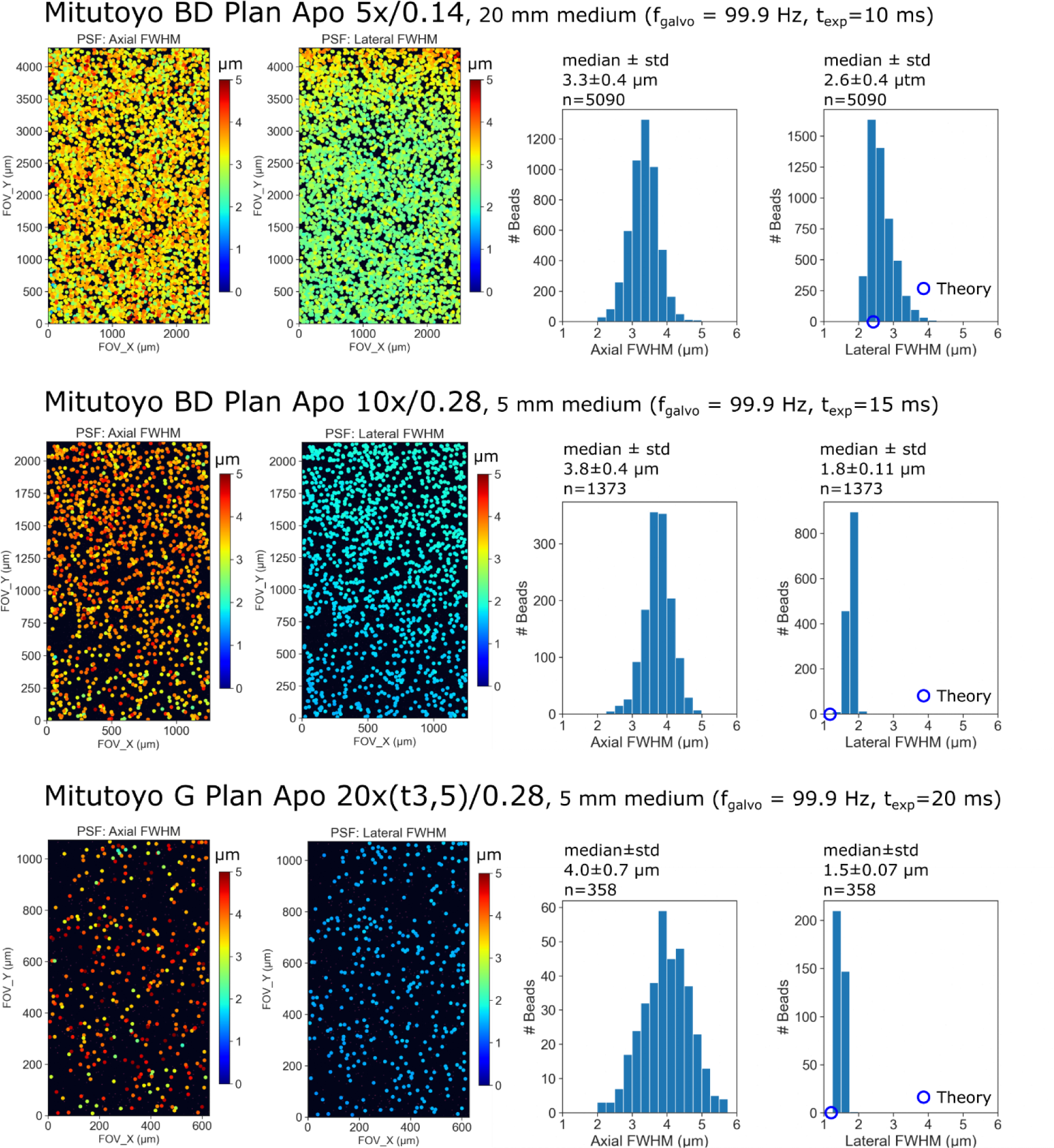
Light-sheet PSF analysis across the field for Mitutoyo Plan Apo detection objectives. PSF was computed from image stack of dense bead sample (200 nm fluorescent beads) embedded in agarose and index-matched with RI=1.52 medium (CUBIC-R+). The overall thickness of high-index medium (glass, oil, and CUBIC-R+, 5-20 mm, RI=1.52) designated as ‘oil’ is indicated. Theoretical values for lateral resolution were calculated as *r*_*lateral*_ = 0.61λ_*det*_/*NA*_*det*_ (assuming ideal imaging conditions and no refractive index mismatch).

**SI Fig. 2.**
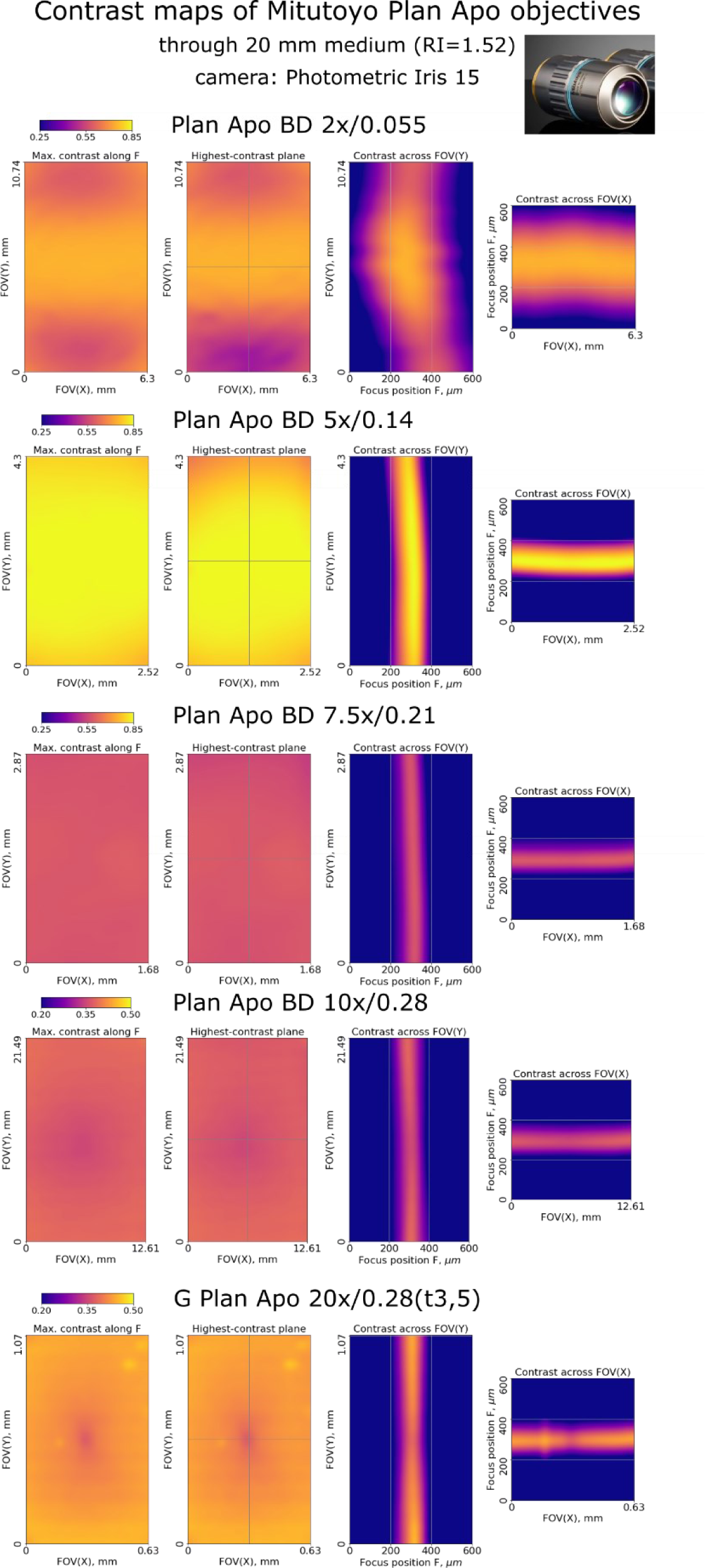
Contrast maps of Mitutoyo Plan Apo objectives. Imaging of a Ronchi ruling was through 20 mm of high-RI medium (RI 1.52). Note that colorbar scale is adjusted to smaller range for 10x and 20x objectives (NA 0.28) to account for spherical aberration effect due to higher NA. The standard tube lens (Mitutoyo MT-1) was used for tests.

**SI Fig. 3.**
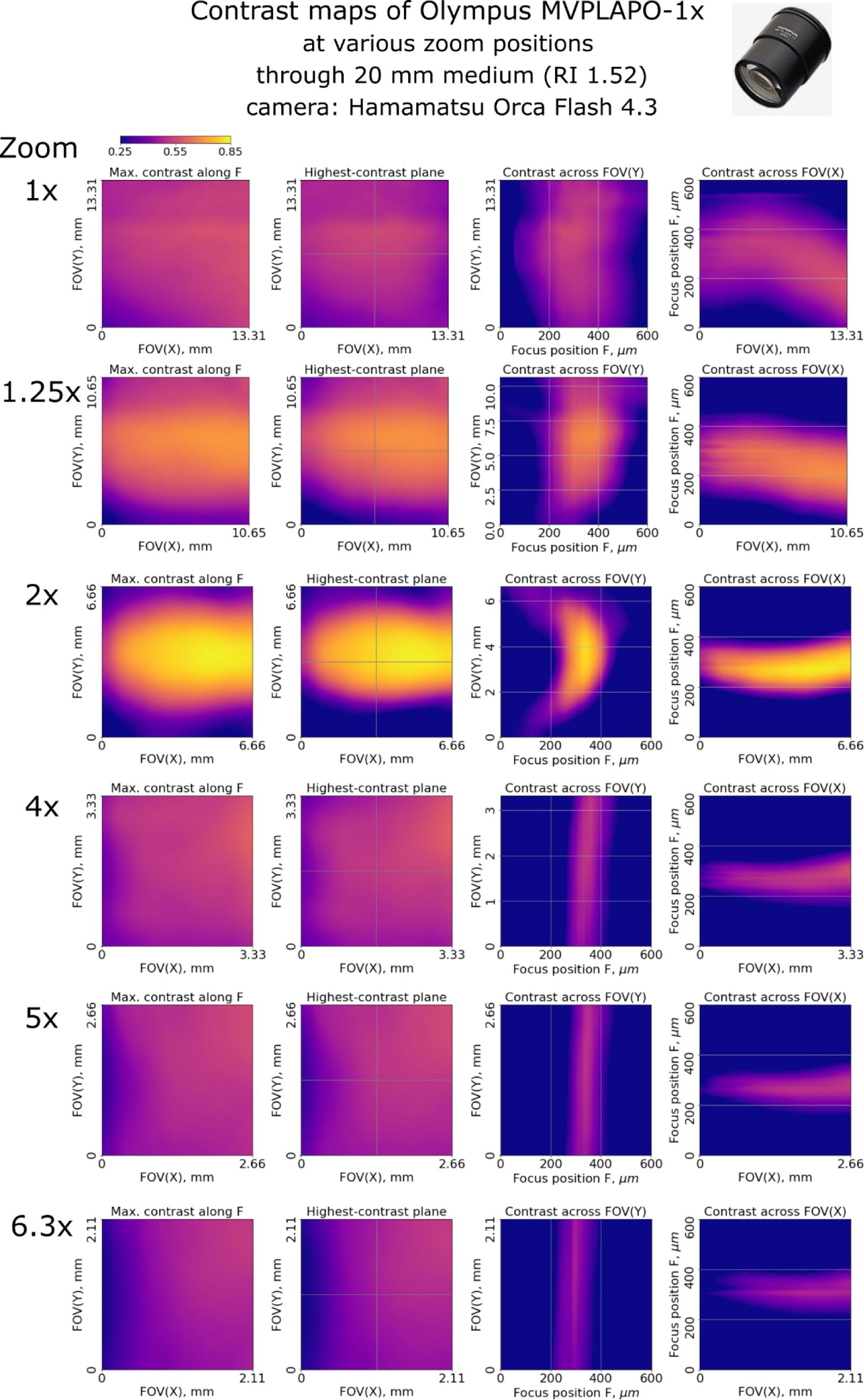
Contrast maps of Olympus MVPLAPO-1x detection objective at various zoom settings. The zoom was set using the Olympus MVX-10 microscope body (mesoSPIM v.5 configuration). Images of the Ronchi ruling were taken through 20 mm of immersion medium (RI=1.52) to mimic the typical imaging conditions.

**SI Fig. 4.**
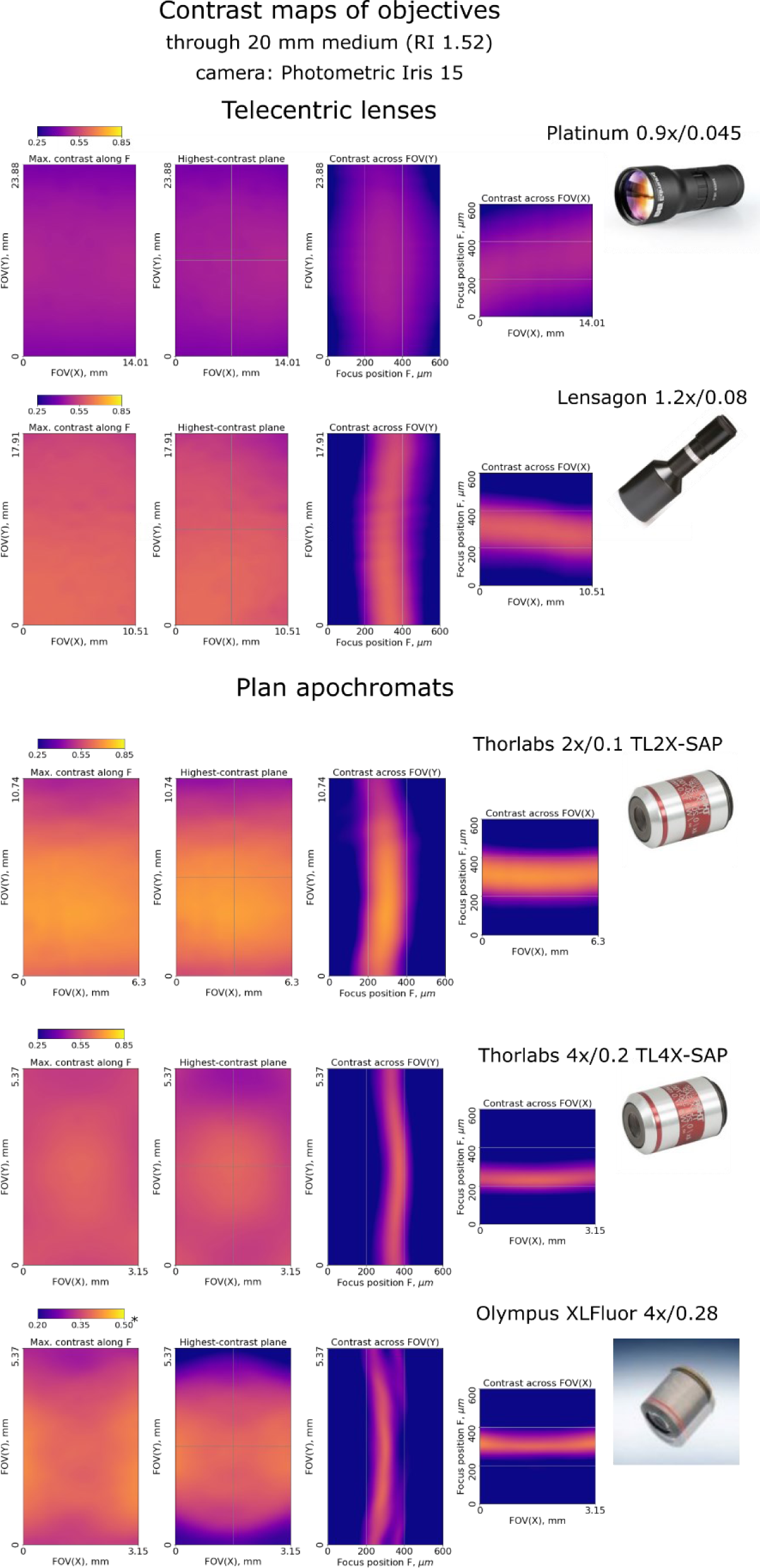
Contrast maps of some other detection objectives. Imaging of Ronchi grating was through 20 mm of high-index medium (RI=1.52). Note that colorbar scale is adjusted to smaller range for the Olympus 4x/0.28 to account for spherical aberration effect at higher NA.

**SI Fig. 5.**
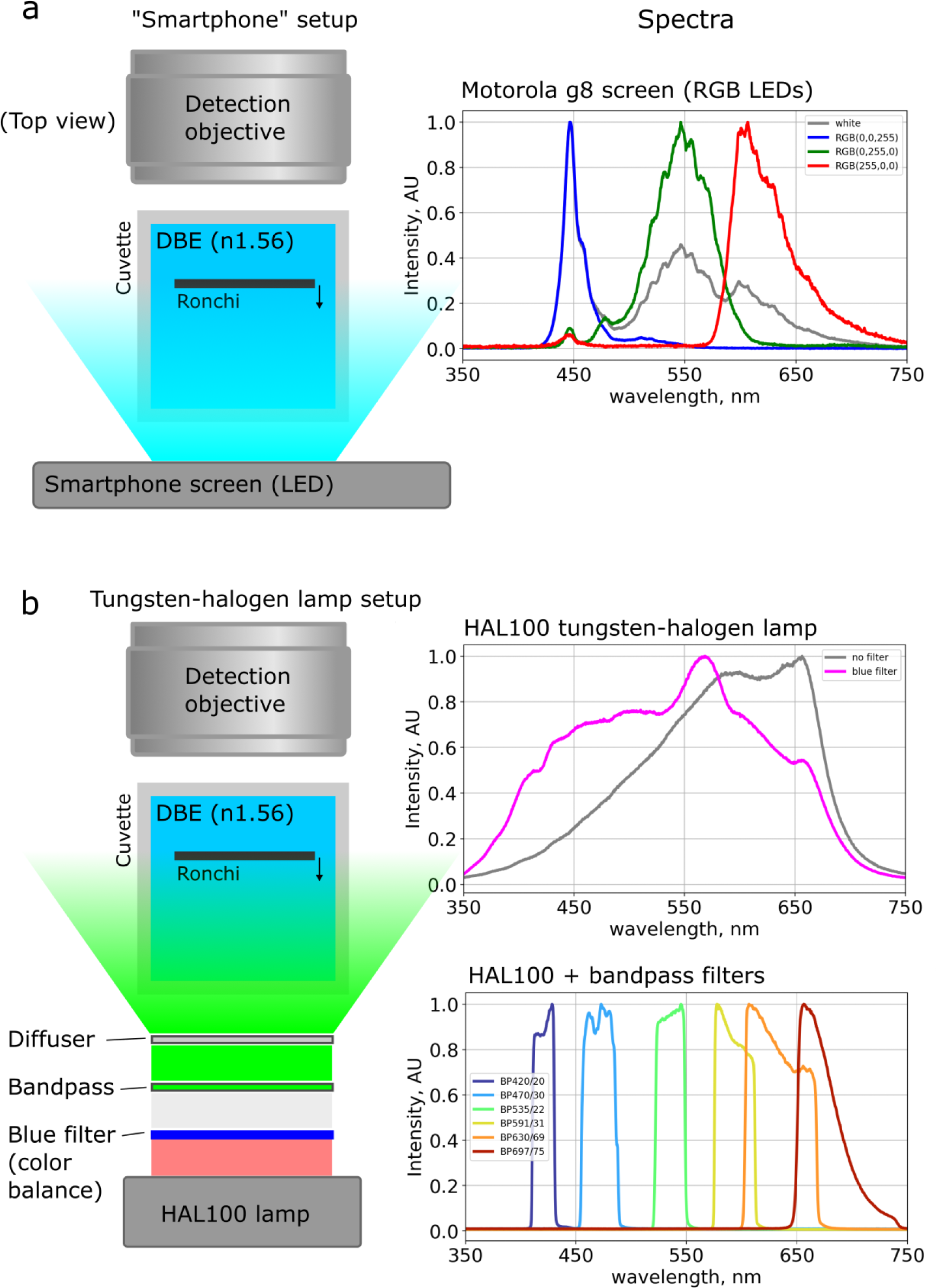
Setups and spectra of the light sources used for contrast map testing. a), The basic illumination setup for objective contrast mapping. Left, top view of the testing setup, with the Ronchi slide immersed in a cuvette with DBE and illuminated by the smartphone screen from behind. Right, – spectra of the smartphone screen with uniform RGB slide (white – (255,255,255), etc). b) The extended illumination setup used for quantification of the chromatic effects, with a broad-spectrum HAL100 tungsten-halogen lamp and bandbass filters. Left: top view of the testing setup. Right: Spectrum of the tungsten-halogen lamp without and with a blue filter for color balance (top) and spectral bands provided by using the 6 bandpass filters (bottom).

**SI Fig. 6.**
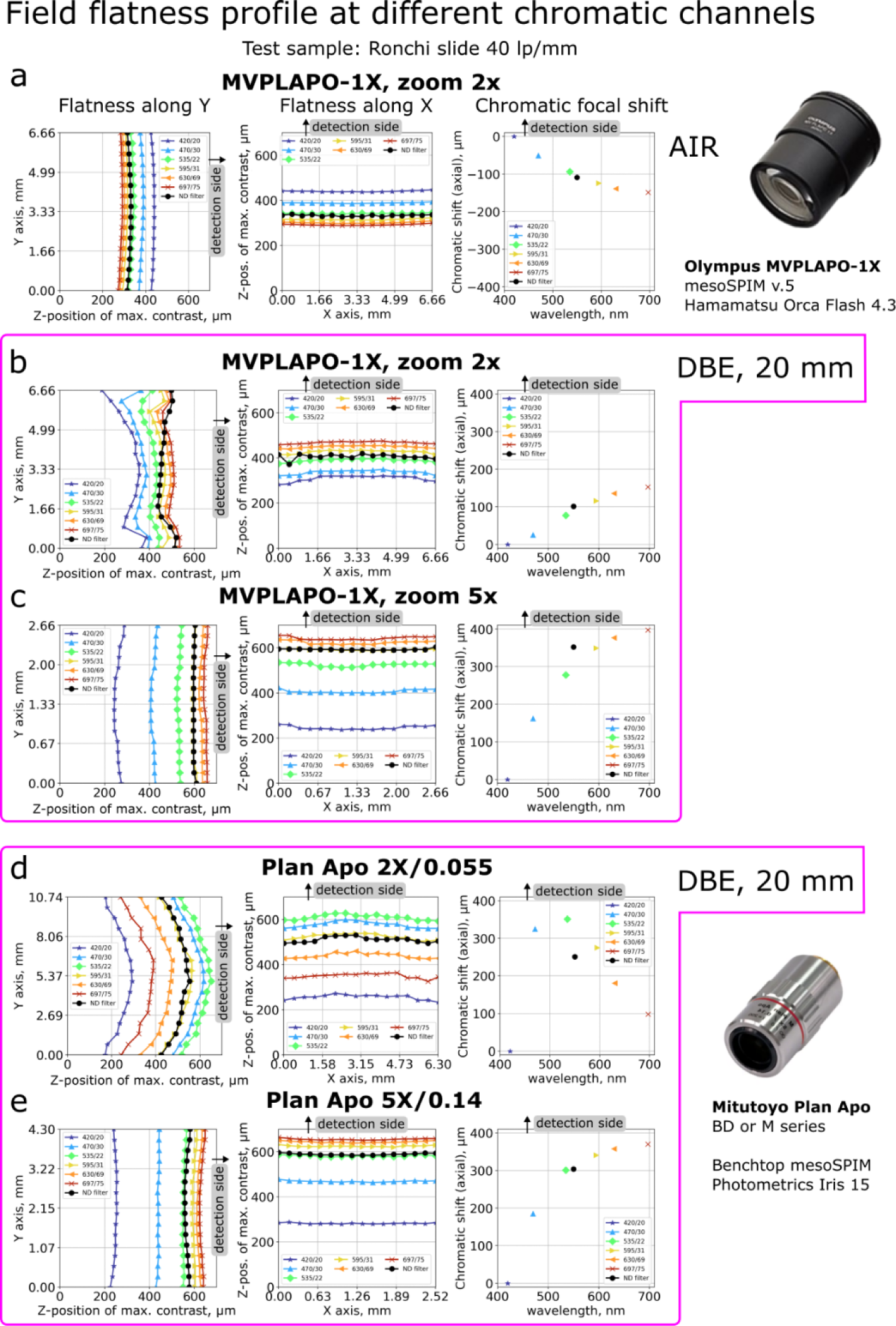
Sections of the best-focus surface at different chromatic channels for select plan apochromat objectives. The test target, Ronchi slide 40 lp/mm, was in the air or immersed in DBE medium (20 mm total, including cuvette glass wall). **a,** Olympus MVPLAPO-1x on MVX-10 body at zoom settings 2x and Ronchi slide in the air; **b**, Ronchi slide immersed in DBE; **c,** zoom setting 5x, Ronchi slide in DBE. **d,** Mitutoyo Plan Apo BD 2x/0.055, Ronchi slide in DBE; **e,** Mitutoyo Plan Apo BD 5x/0.14, Ronchi slide in DBE. Note that Mitutoyo objectives were tested with a Photometrics Iris 15 camera, which has sensor significantly larger in Y axis and slightly smaller in X axis than Hamamatsu Orca Flash 4.3 (Fig. 1d of the main text). Measurements for Mitutoyo BD also applies to equivalent objectives in M series. In each row, left and middle panels have z-position zero chosen arbitrary, to cover the full range of profiles in all channels; the right panel shows profile offsets relative to the “420/20 nm filter” measurement.

**SI Fig. 7.**
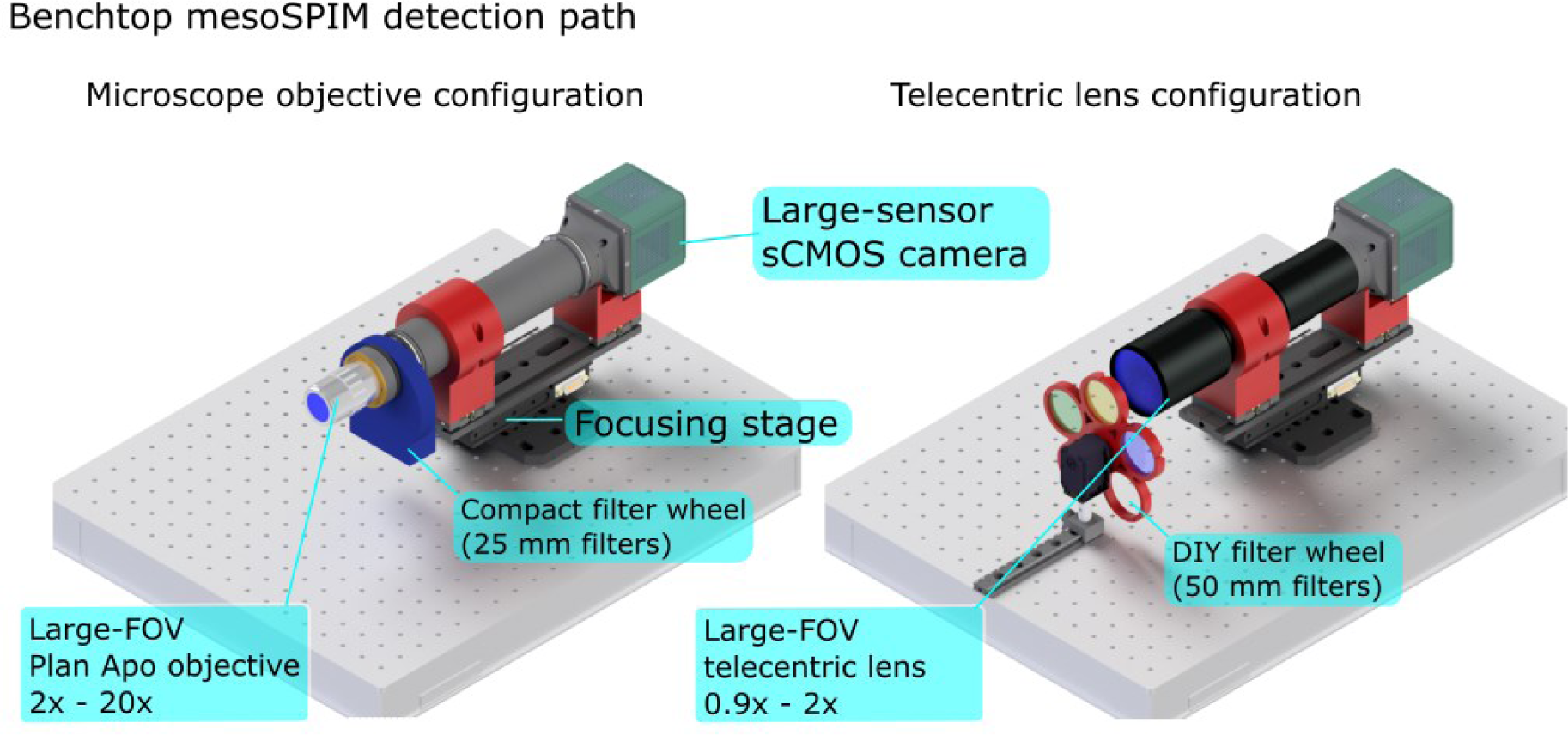
The Benchtop mesoSPIM detection path design. CAD drawing of detection arm mechanical design. Two main options are shown: microscope objective (2x – 20x) vs telecentric lens (0.5 – 2x). Custom-made (3D-printed) parts are red.

**SI Figure 8.**
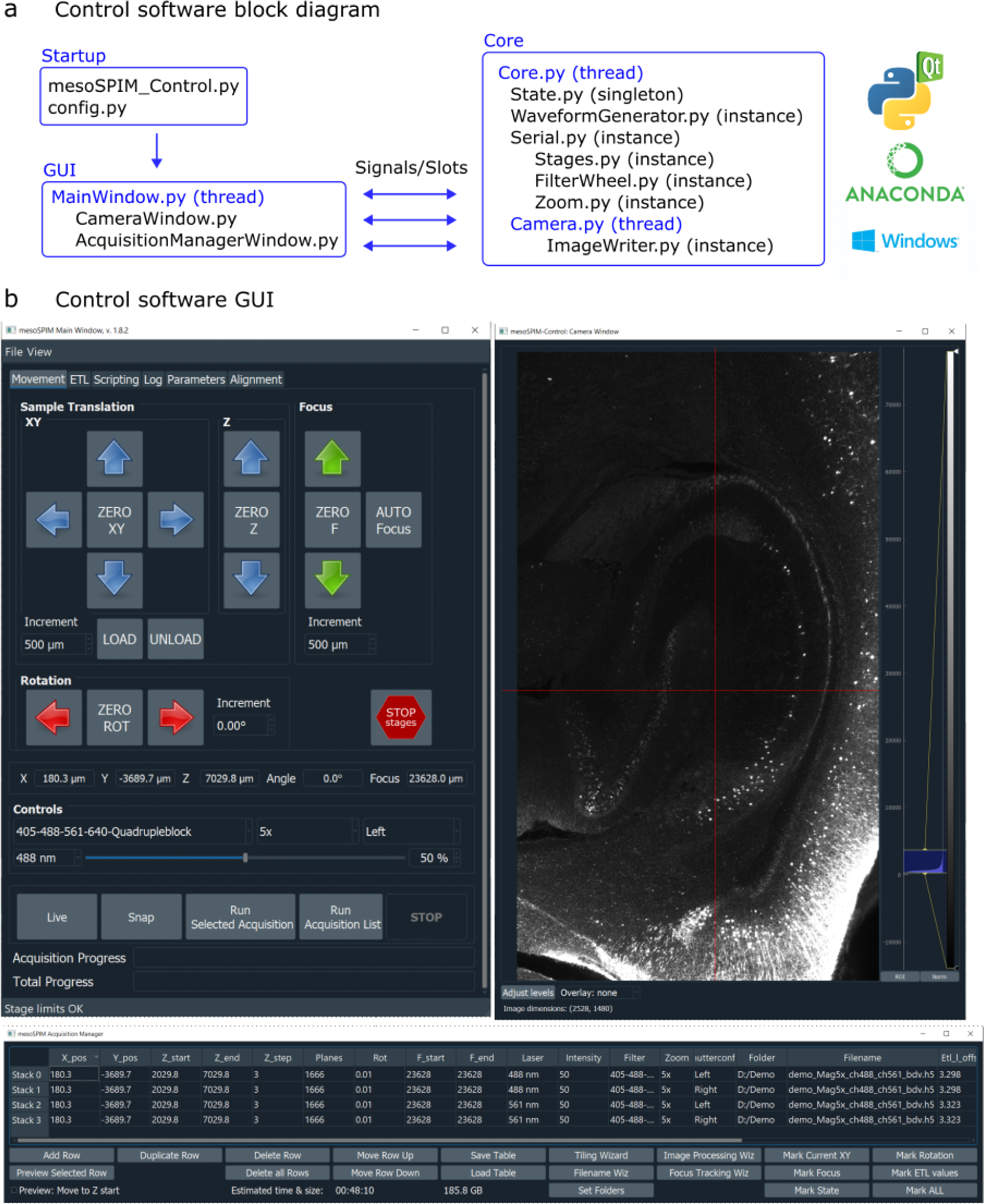
The mesoSPIM control software. **a,**Block diagram: startup file mesoSPIM_Control.py launches system-specific parameters from the config file, creates GUI windows, and initializes core classes and threads. Communication between threads and class instances is managed by PyQt signal/slot mechanisms. Python interpreter and dependencies are managed by Anaconda under Windows 10, 64-bit. **b,** The three main GUI windows, in dark scheme: Main Window with the control parameters, Camera Window with live view of the current imaging plane, and Acquisition Manager which contains all required parameters for each stack/view acquisition.

**SI Fig. 9:**
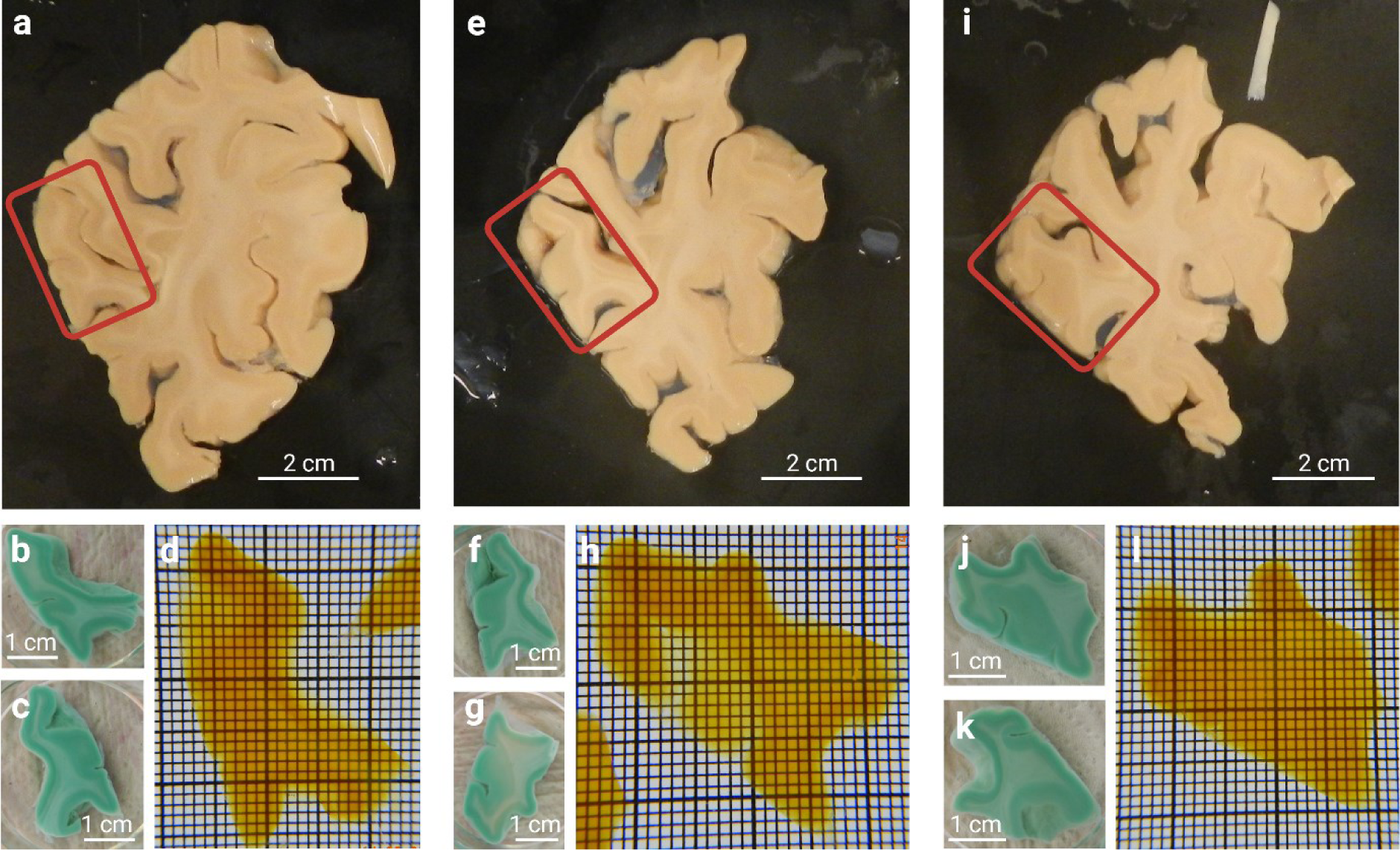
Histological processing of Occipital lobe 1 samples with MASH-MB. Whole occipital lobe slices shown with the anterior side facing up (a, e, and i; sequence of slices from anterior to posterior) were blocked around the macroscopically visible V1/V2 border (red boxes). Blocked samples, after labelling with methylene blue, show distinct contrast between grey and white matter (b, f, and j: anterior side of the samples; c, g, and k: posterior side of the samples). All samples are well cleared and become highly transparent, even in the white matter, after immersion in ECi (**d**, **h**, and **l**). In **d**, **h**, **l**, the grid is 1×1 mm (smallest squares).

**SI Fig. 10:**
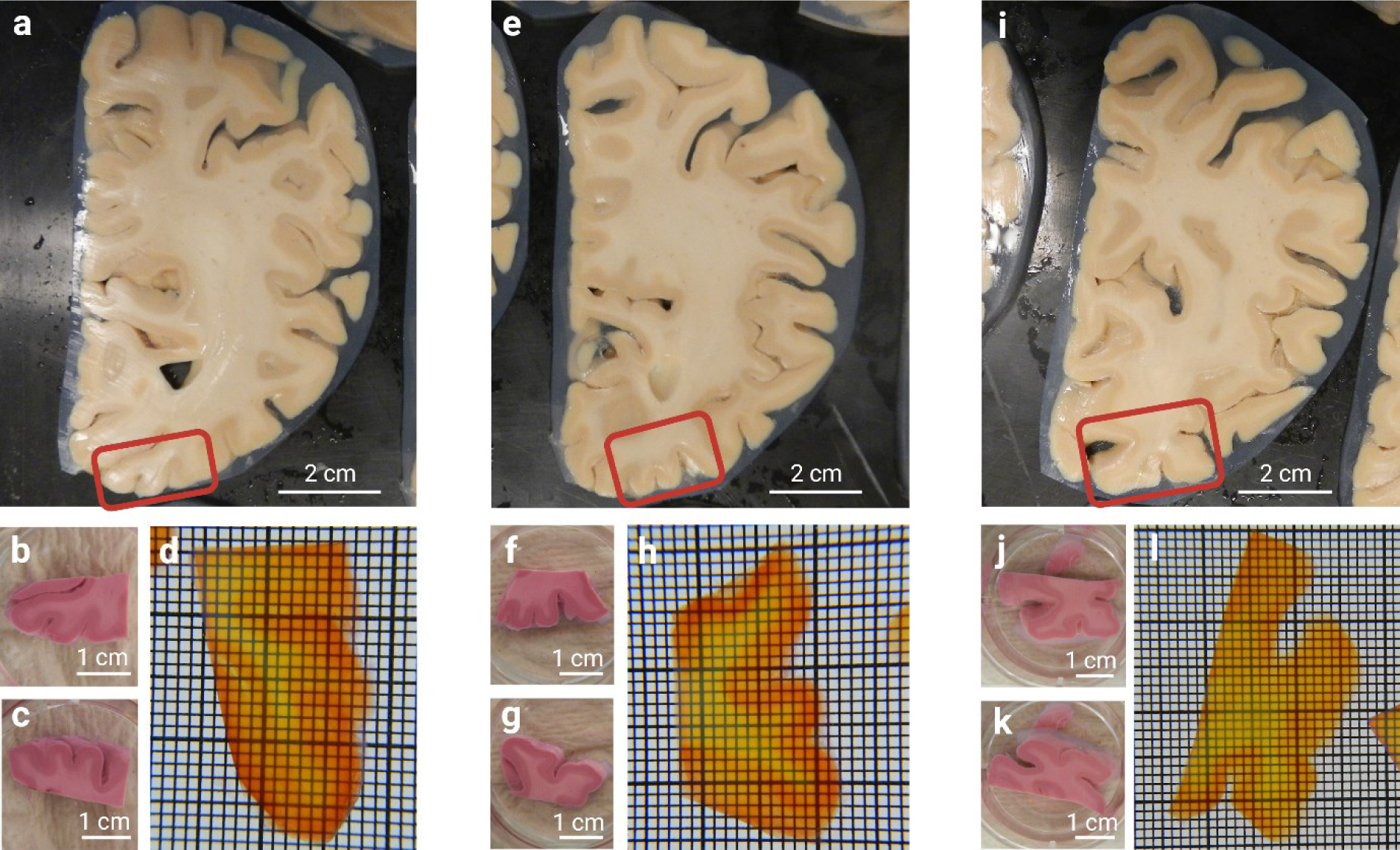
Histological processing of Occipital lobe 2 samples with MASH-NR. Whole occipital lobe slices shown with the anterior side facing up (a, e, and i; sequence of slices from anterior to posterior) were blocked around the macroscopically visible V1/V2 border (red boxes). Blocked samples, after labelling with neutral red, show distinct contrast between grey and white matter (b, f, and j: anterior side of the samples; c, g, and k: posterior side of the samples). All samples are well cleared and become highly transparent, even in the white matter, after immersion in ECi (**d**, **h**, and **l**). In **d**, **h**, **l**, the grid is 1×1 mm (smallest squares).

**SI Fig 11.**
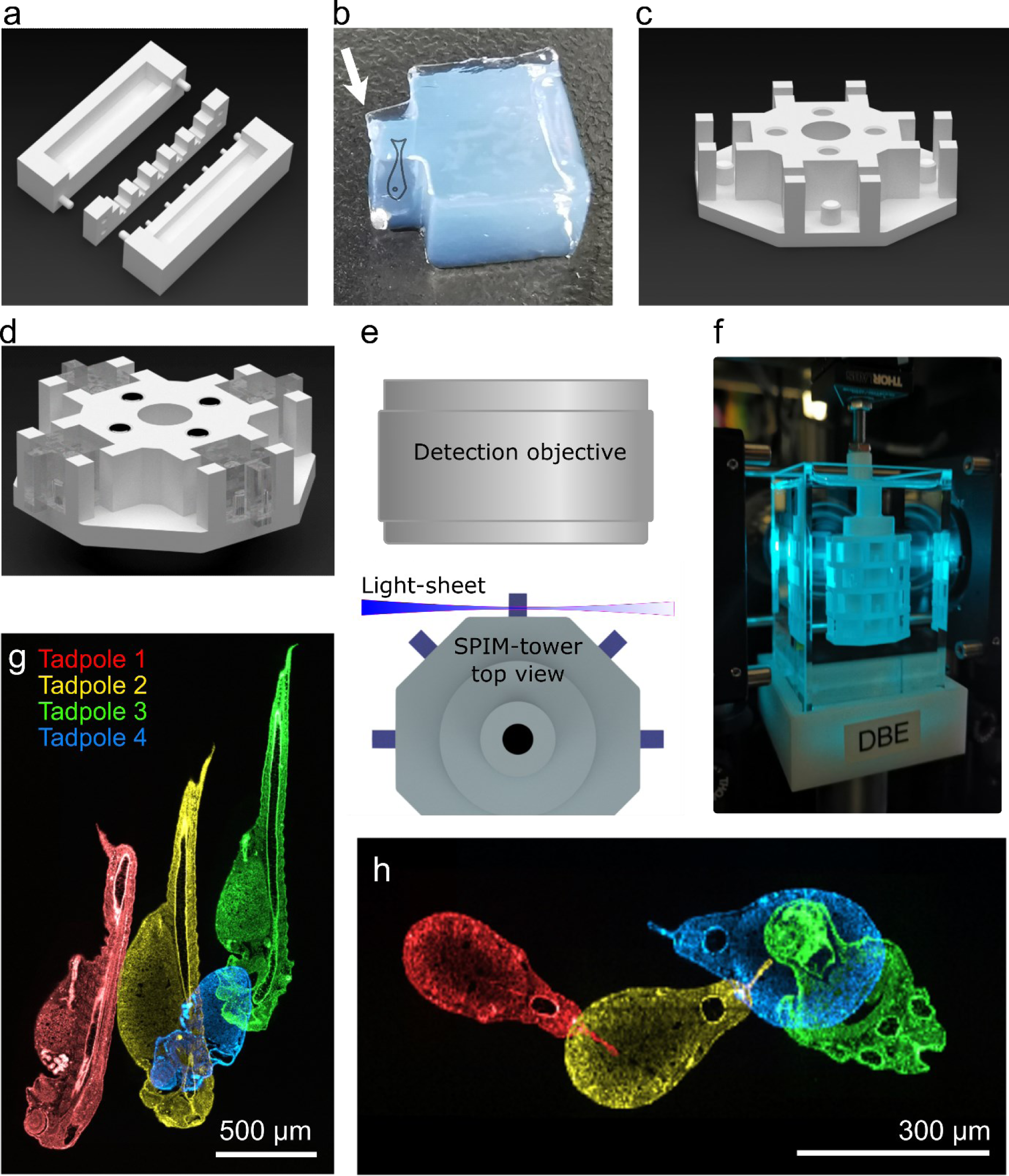
The *SPIM-tower* sample holder and *SPIM-mold*: a standardized high-throughput imaging on the mesoSPIM system. **a,** CAD model of the *SPIM-mold* for reproduceable embedding of biological samples in agarose blocks. The mold has been optimized to allow for the positioning of 6 *X. tropicalis* tadpoles simultaneously in a standardized way. A pin on the *SPIM-mold* generates a negative hole for future mounting into the *SPIM-tower*. **b,** The resulting agar block with embedded sample (tadpole, position indicated by cartoon drawing). The larger part of the block is for sample mounting into the tower, and the smaller part contains the sample (arrow) and protrudes from the tower, like a small balcony. This design facilitates access of the light-sheet to the sample. **c,** One section (level) of the *SPIM-tower* holder, with 4 holes in the center for push-magnets, and 4 sample slots positioned for mounting agar blocks. Each slot has a mounting pin to fit securely into the hole in the agar block. This allows consistent and reproducible mounting of samples into the *SPIM-tower*. **d,** Four agar blocks containing samples, mounted onto one level of the *SPIM-tower*. The push-magnets allow stacking multiple levels to form a tower. **e,** Top view of the *SPIM-tower* with light-sheet illuminating the sample, and the detection objective shown. Consecutive levels are rotated by 45° from each other to avoid samples from different levels interfering with each other. **f,** The *SPIM-tower* in the Benchtop mesoSPIM during imaging. Each sample can be accessed by 90° rotation within the same level and Y translation (7.5 mm) + 45° rotation from one level to the next. **g,h** Visualization of the deviation in position of 4 stage-37 tadpoles (pseudo-colored) on one SPIM-tower level in the X-Y (**g**) and XZ (**h**) axes. Due to the *SPIM-mold* design and standardized agarose blocks, the X, Y deviations are within 1 mm and Z within 0.3 mm.

**SI Fig 12.**
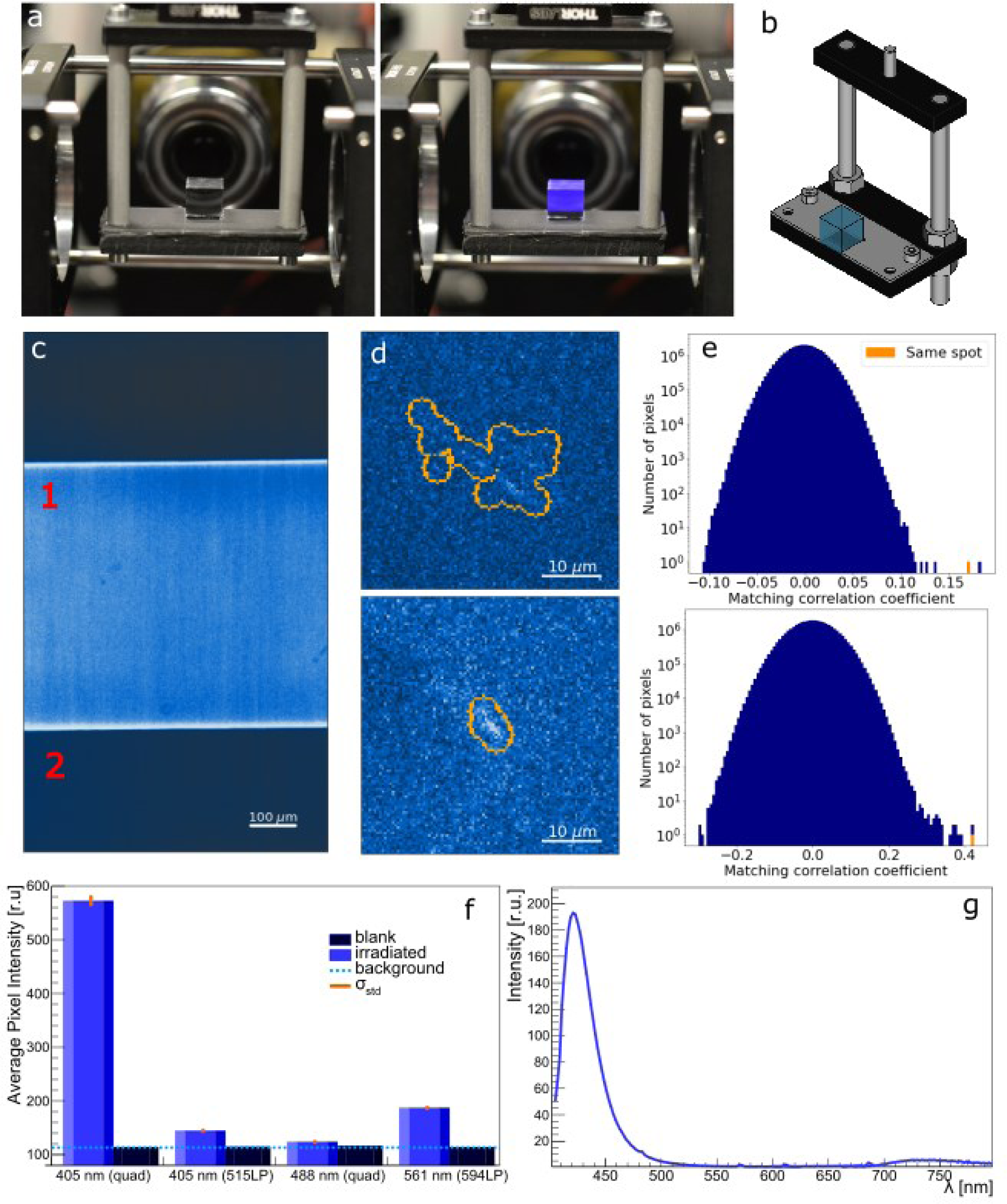
Imaging color centers in CaF2 crystal in the Benchtop mesoSPIM. The crystal was irradiated with gamma rays at 5 MRad, and imaged using Mitutoyo 20x/0.28 (t3,5) objective and 405nm excitation, without immersion. **a,** The gamma-irradiated CaF2 crystal is highly fluorescent in the blue spectrum when illuminated with 405-nm laser (left panel: laser off, right panel: laser on). **b**, crystal sample holder design for cubic crystals, no immersion (imaging in air). **c,** Light-sheet illuminated area (bright band, ROI #1) is compared to non-illuminated areas (two dark bands, with ROI#2) for a irradiated CaF2 crystal. **d**, Clusters of high-intensity pixels found in the scans of a CaF2 crystal using the algorithm described in **Sl Note 2**. **e,** Matching correlation coefficients of groups of N pixels in repeated scans. The bins in orange show the matching correlation coefficients of the same group of pixels within the contours shown in the images on the left side of each graph. See **Sl Note 2** for a detailed description. **f,** Quantification of crystal fluorescence from blank and irradiated (100 kRad) CaF2 crystals imaged at 1x with different excitation laser wavelengths and emission filters. The background line represents the camera noise measured when the laser is off. The small error bars in orange show the standard deviation of the average pixel intensities from 350 z-planes. The solid bars show the mean value of the distribution. **g,** fluorescence spectra from the same irradiated CaF2 crystal measured in response to 400 ± 10 nm excitation light.

**SI Fig 13.**
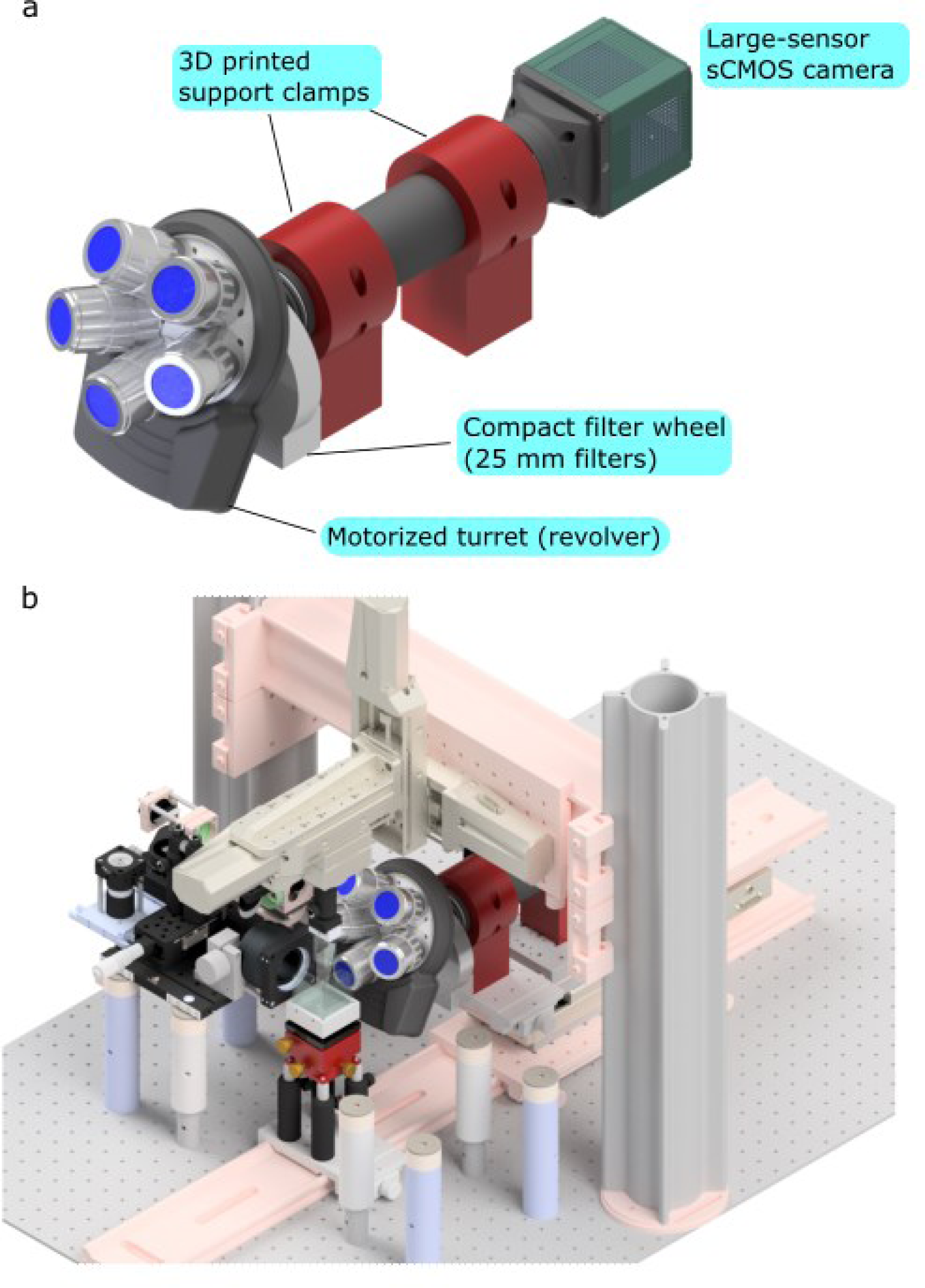
CAD model of the new detection arm for mesoSPIM v4-5 upgrade. **a,**The detection arm elements. The height of support allows drop-in replacement into the existing detection path of mesoSPIM v4-5; **b,** The new detection arm in mesoSPIM v.5. The right excitation arm is invisible for presentation purposes.

**SI Fig. 14.**
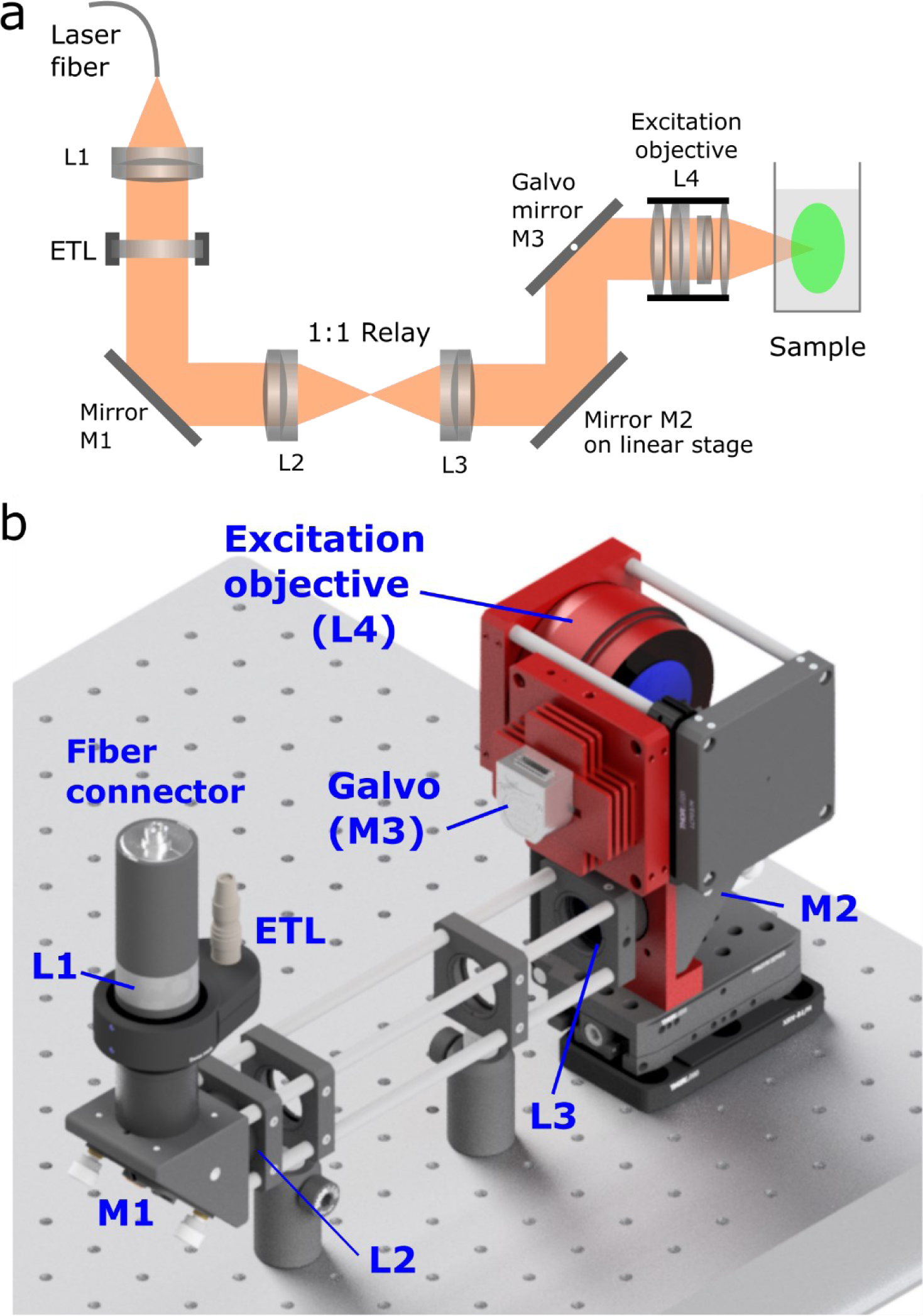
The Benchtop mesoSPIM excitation arm. **a**, Diagram of the laser beam propagation and optical components. **b**, Mechanical implementation of the left excitation arm (view from behind). Custom-made or modified parts are color-coded in red (mirror bracket, galvo mount, galvo heat sink, Nikon 50 mm f/1.4 G and its mount). All parts and details are available at the github repository.

**SI Fig. 15.**
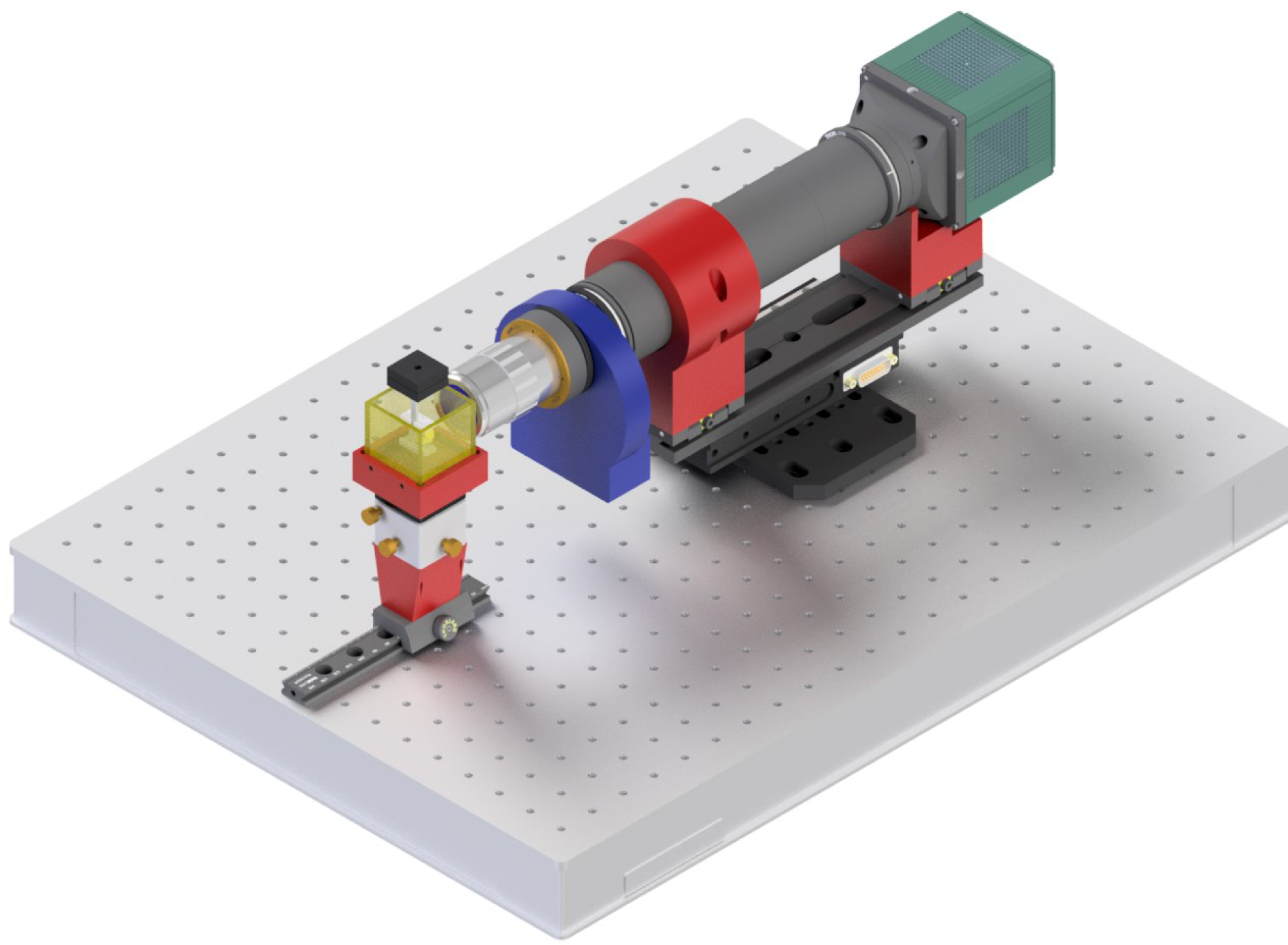
Immersion chamber in the detection path of Benchtop mesoSPIM.

**SI Fig. 16.**
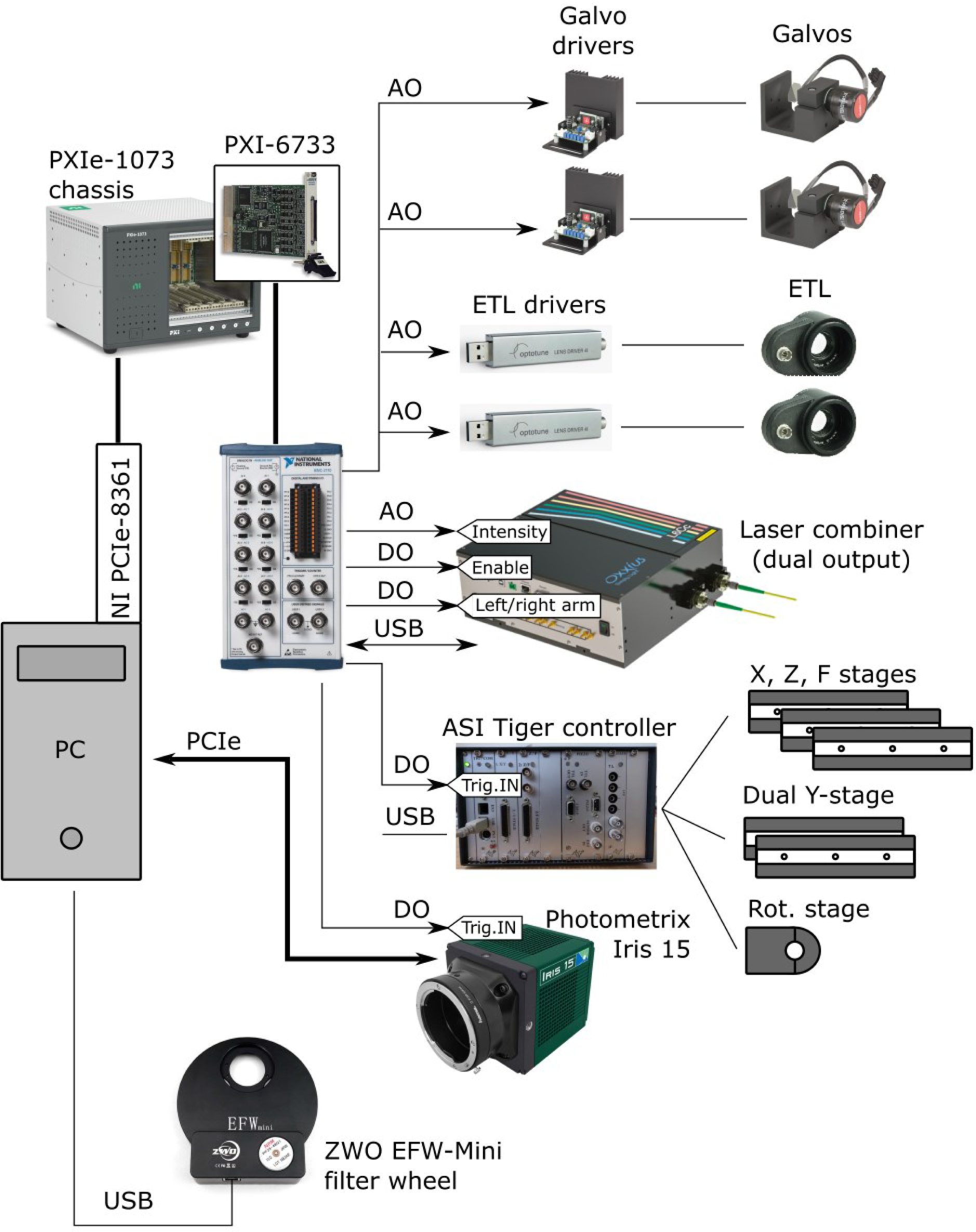
Electronics block diagram of Benchtop mesoSPIM.

**SI Notes Fig. 1:**
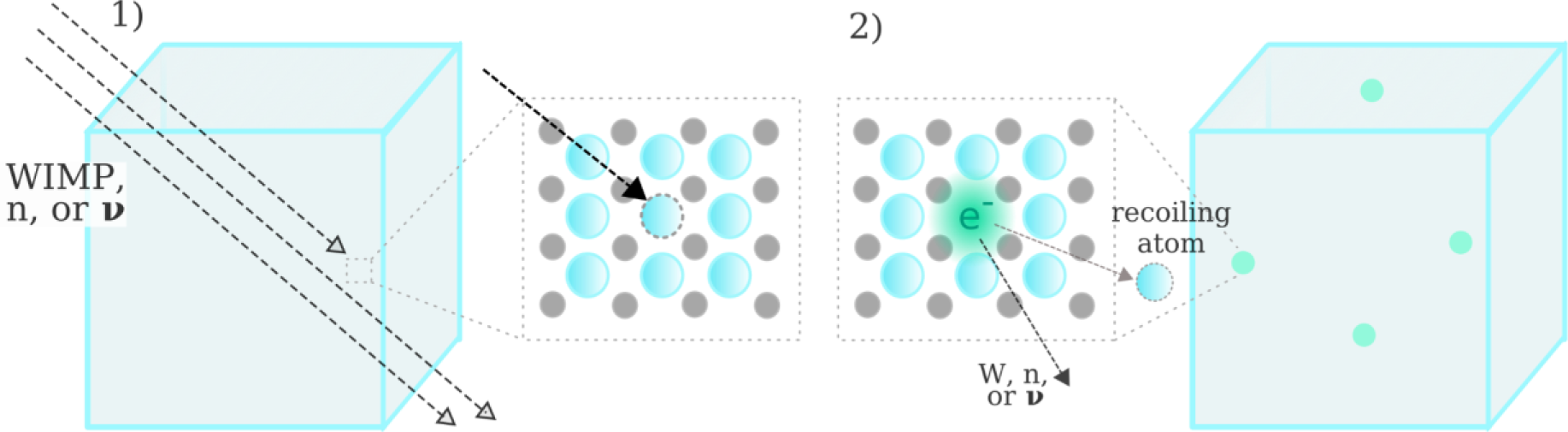
1) A transparent crystal is exposed to a flux from particles: dark matter (Weakly Interacting Massive Particle, WIMP), neutrons (n) or neutrinos (*v*). If one of these particles interacts (collides) with a nucleus, the atom can recoil and be permanently displaced from its lattice position. In the case of an anion, the vacancy can be occupied by an electron, forming an F-center type of color center^14^. 2) This center appears as a fluorescent point in a SPIM image of the crystal. For more massive/interacting particles, such as cosmic-rays, multiple atoms can be displaced forming a *track* of color centers. For gamma-rays, no atom displacement is expected, but ionization and consequent charge redistribution can also create color centers of different types, as described in Ref^14^. After exposure, the color centers are imaged with SPIM and their position and intensity are recorded.

**SI Notes Fig. 2.**
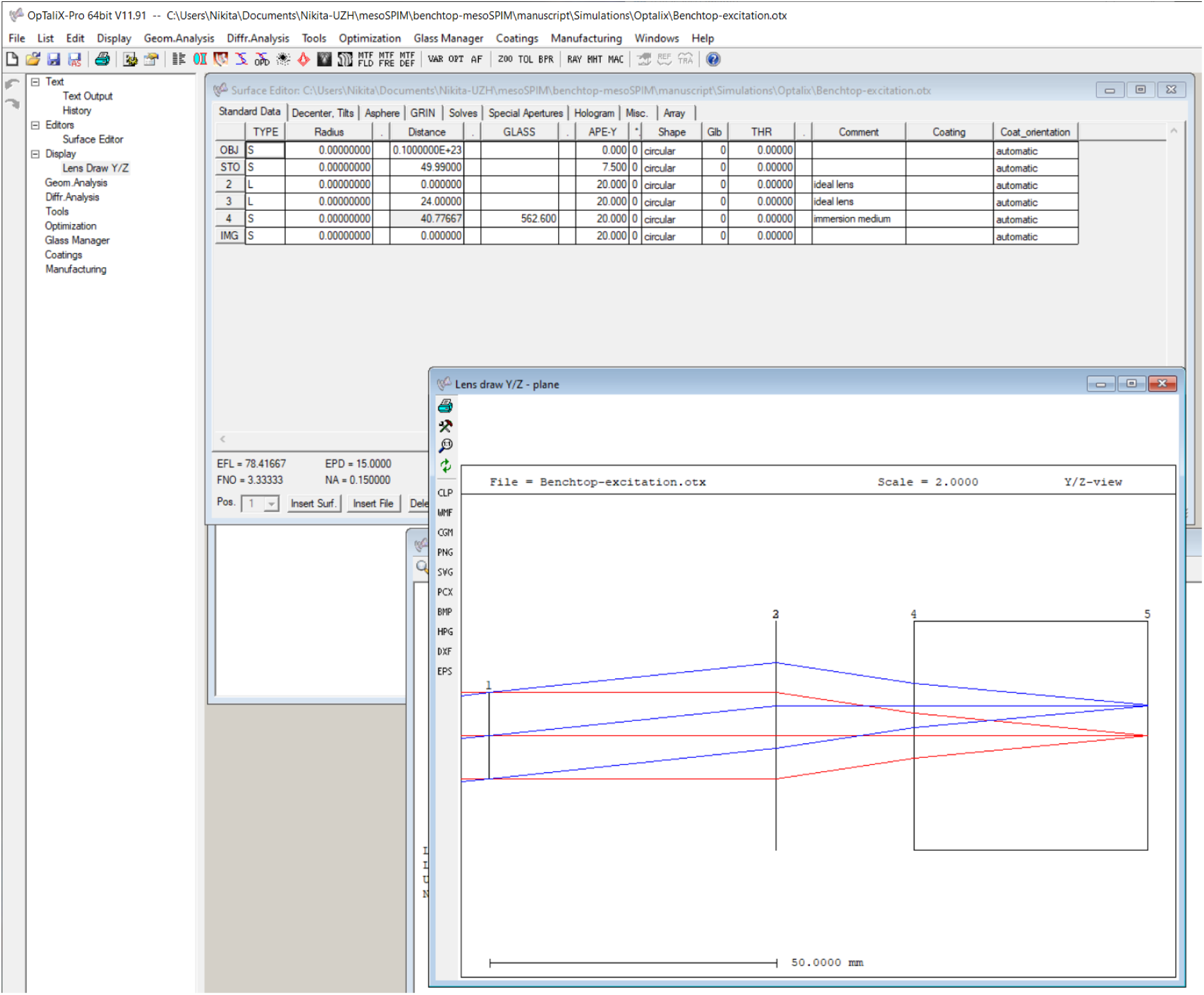
Simplified model of the excitation objective and immersion chamber. The surface #4 is where the immersion medium (refractive index 1.562) starts. Surface #5 is the plane where beam profile PSF is evaluated. Apodization was set to 0.135 to mimic the Gaussian beam.

**SI Notes Fig. 3.**
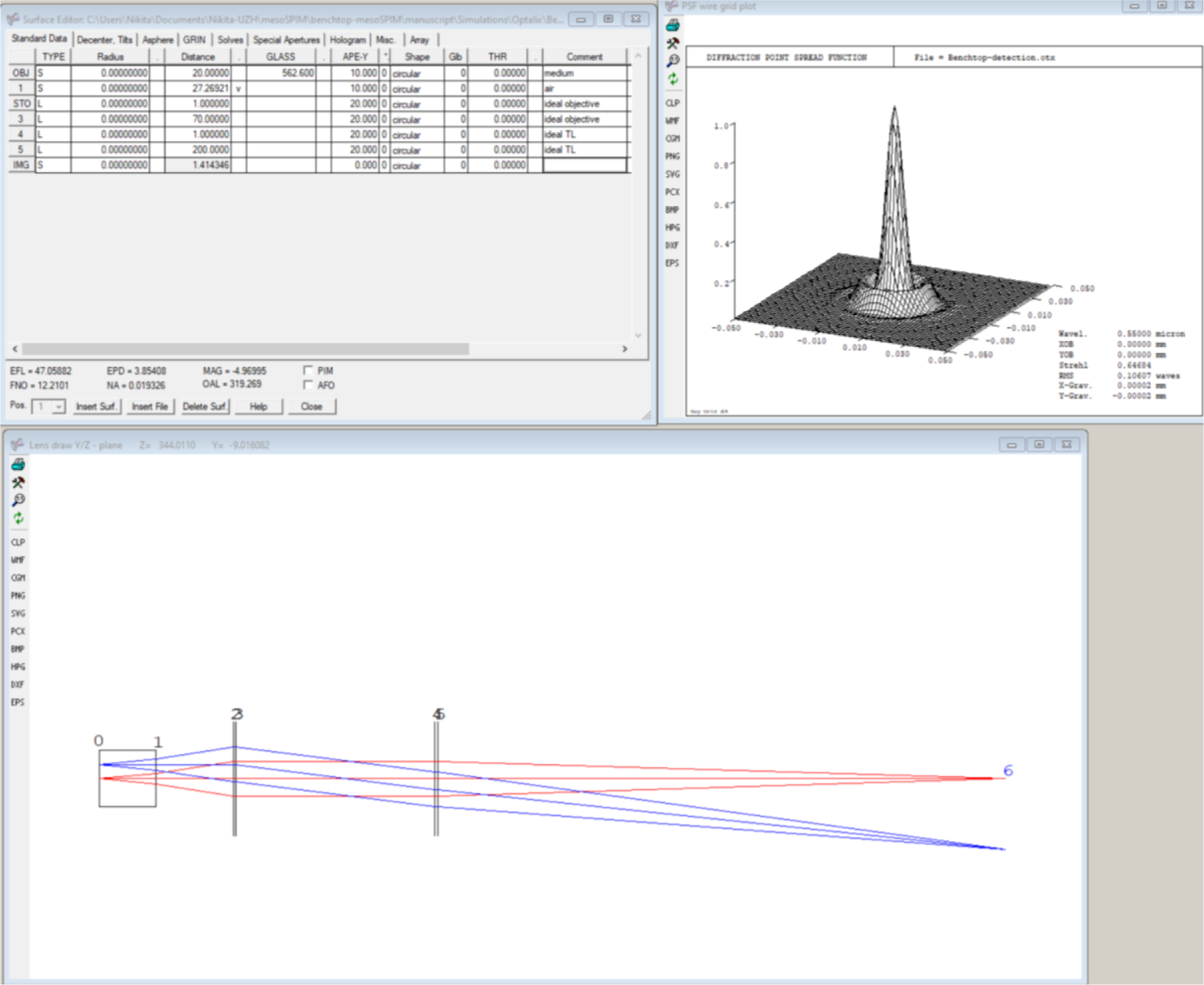
Simplified model of the detection path (immersion chamber, 5x objective, and a tube lens) in OptalixPro software.

## Supplementary tables

**SI Table 1.** Properties of microscope objectives and telecentric lenses computed from their contrast maps.

| Objective | Field sag<br>over FOV_Y<br>(long axis),<br>$\mu\text{m}$ | Field sag<br>over FOV_X<br>(short axis),<br>$\mu\text{m}$ | Max contrast in<br>the best-focus<br>plane | Ave contrast in<br>best-focus plane | Min contrast in<br>the best-focus<br>plane |
| --- | --- | --- | --- | --- | --- |
| PlatinumTL™ 0.9x Telecentric Lens | 21 | 20 | 0.49 | 0.45 | 0.37 |
| Lensagon T25M-12-155I Telecentric Lens | 42 | 5 | 0.63 | 0.58 | 0.45 |
| Mitutoyo BD Plan Apo 2x/0.055 | 90 | 22 | 0.75 | 0.65 | 0.45 |
| Mitutoyo BD Plan Apo 5x/0.14 | 18 | 10 | **0.86** | **0.82** | **0.65** |
| Mitutoyo BD Plan Apo 7.5/0.21 | **4** | **4** | 0.60 | 0.58 | 0.53 |
| Mitutoyo BD Plan Apo 10/0.28 | < 1 | **4** | 0.39 | 0.37 | 0.35 |
| Mitutoyo G Plan Apo 20x/0.28(t3.5) | **1** | **2** | 0.48 | 0.43 | 0.37 |
| Olympus MVPLAPO-1x, zoom 1x | 50 | 70 | 0.57 | 0.48 | 0.34 |
| Olympus MVPLAPO-1x, zoom 1.25x | 22 | 11 | 0.70 | 0.54 | 0.14 |
| Olympus MVPLAPO-1x, zoom 2x | 175 | 36 | 0.84 | 0.52 | 0.11 |
| Olympus MVPLAPO-1x, zoom 4x | 28 | 20 | 0.60 | 0.49 | 0.35 |
| Olympus MVPLAPO-1x, zoom 5x | 19 | 15 | 0.55 | 0.47 | 0.29 |
| Olympus MVPLAPO-1x, zoom 6.3x | 13 | 12 | 0.53 | 0.42 | 0.25 |
| Thorlabs Super Apochromatic 2x/0.1 TL2X-SAP | 51 | 13 | 0.72 | 0.62 | 0.43 |
| Thorlabs Super Apochromatic 4x/0.2 TL2X-SAP | 39 | 11 | 0.62 | 0.55 | 0.44 |
| Olympus XLFluor 4x/0.28 | 26 | 15 | 0.41 | 0.33 | 0.18 |

**SI Table 2.** Currently recommended objectives for Benchtop mesoSPIM.

| Model | Mag/NA | WD, mm | Resolution, $\mu\text{m}$ | Max image circle (diagonal), mm | Approx cost, USD | Notes |
| --- | --- | --- | --- | --- | --- | --- |
| **PlatinumTL™ 0.9x Telecentric Lens** | 0.9x/0.045 | 111 | 5.5 | 28.7 | 2500 | Telecentric lens* |
| **Lensagon T25M-12-155l** | 1.2x/0.08 | 155 | 4.2 | 32 | 6700 | Telecentric lens* |
| **Mitutoyo (M or BD) Plan Apo** | 2x/0.055 | 34 | 5 | 30 | 1100 | Mitutoyo lens**. Field flatness acceptable. |
| **Mitutoyo (M or BD) Plan Apo** | 5x/0.14 | 34 | 2 | 30 | 900 | Mitutoyo lens**. Field flatness excellent. Spherical aberration minimal even in thick samples. |
| **Mitutoyo (M or BD) Plan Apo** | 7.5x/0.21 | 34 | 1.3 | 30 | 1800 | Mitutoyo lens**. Field flatness excellent. Spherical aberration acceptable for samples up to 10 mm thick. |
| **Mitutoyo (M or BD) Plan Apo** | 10x/0.28 | 34 | 1.0 | 30 | 1200 | Mitutoyo lens**. Field flatness excellent. Spherical aberration acceptable for samples up to 5 mm thick. |
| **Mitutoyo G Plan Apo 20x(t3,5)** | 20x/0.28 | 29.42 | 1.0 | 30 | 3600 | Mitutoyo lens**. Field flatness excellent. Spherical aberration corrected for samples up to 3.5 mm thick (n1.52). |

\* F-mount directly to the camera. Filter wheel (2"-filters) in front of the lens. Galvos require active (fan) cooling.
\*\* Requires Mitutoyo MT-1 (f=200m) or similar large-FOV tube lens. Filter wheel (1"-filters) behind the objective (infinity space). The BD version has larger diameter and mounting thread and requires 3D printed screen ring.

**SI Table 3.**
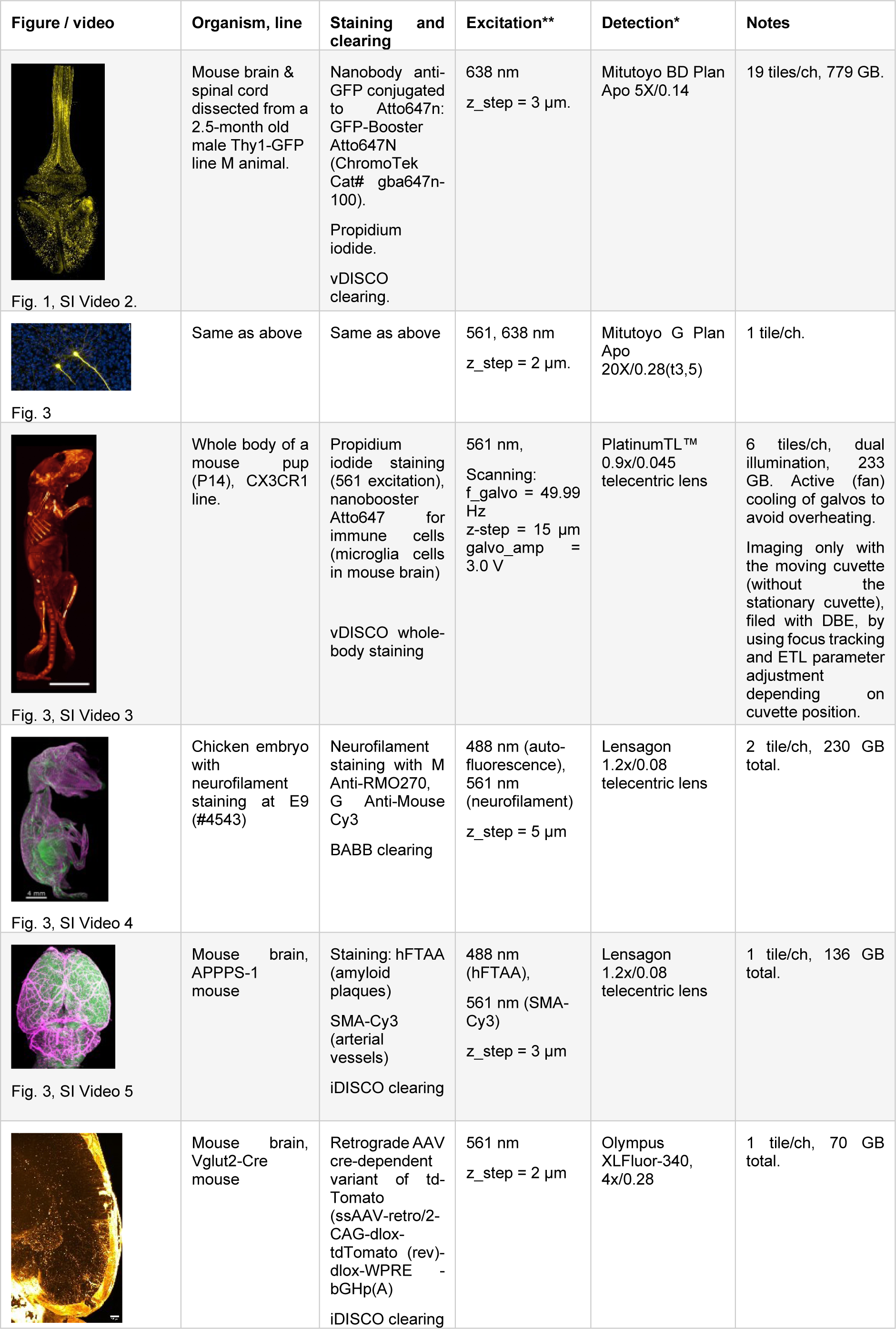
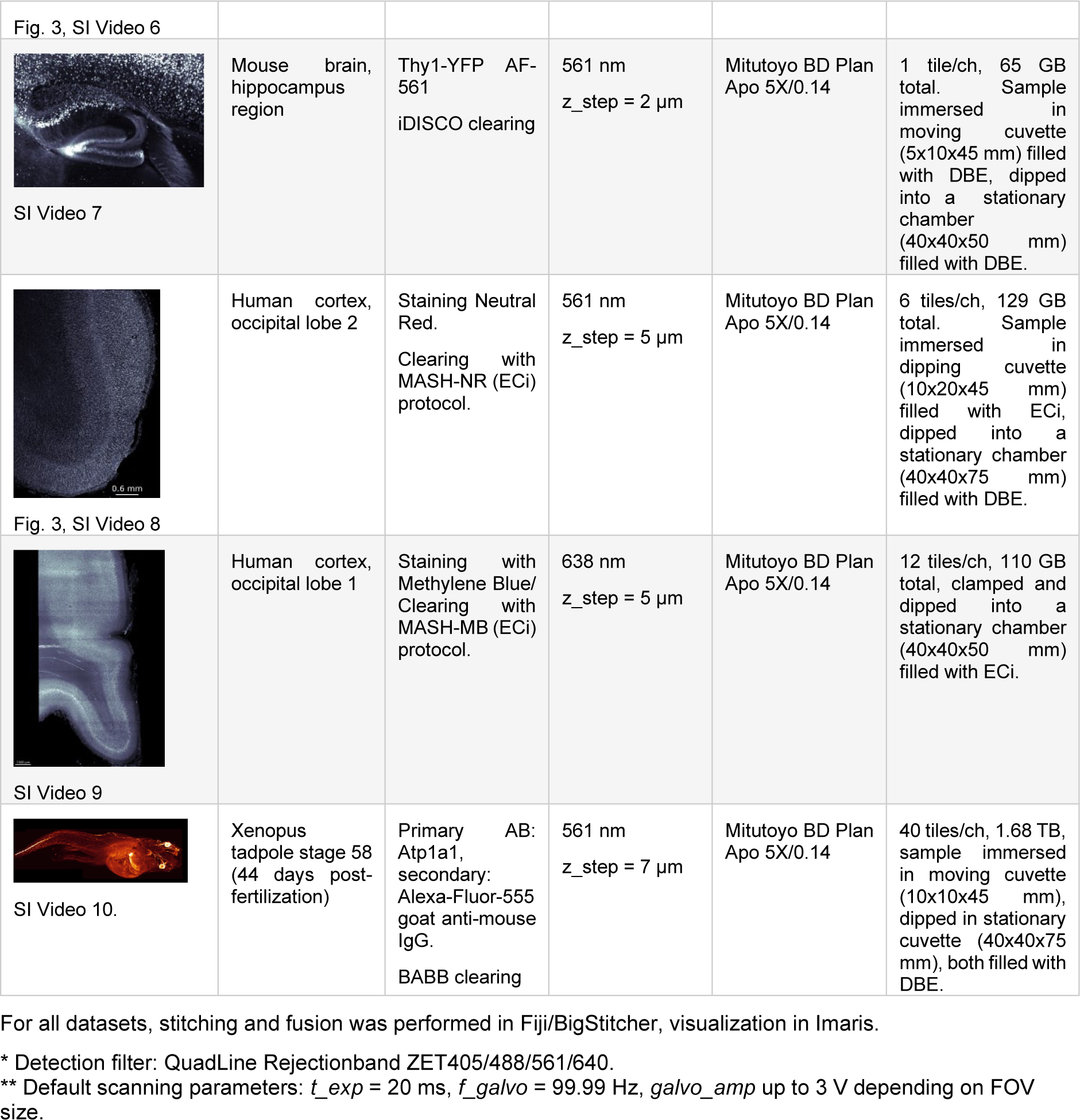
Samples, labelling, clearing, and acquisition parameters.

| Figure / video | Organism, line | Staining and clearing | Excitation** | Detection* | Notes |
| --- | --- | --- | --- | --- | --- |
| Fig. 1, SI Video 2. | Mouse brain & spinal cord dissected from a 2.5-month old male Thy1-GFP line M animal. | Nanobody anti-GFP conjugated to Atto647n: GFP-Booster Atto647N (ChromoTek Cat# gba647n-100).<br>Propidium iodide.<br>vDISCO clearing. | 638 nm<br>z_step = 3 μm. | Mitutoyo BD Plan Apo 5X/0.14 | 19 tiles/ch, 779 GB. |
| Fig. 3 | Same as above | Same as above | 561, 638 nm<br>z_step = 2 μm. | Mitutoyo G Plan Apo 20X/0.28(t3,5) | 1 tile/ch. |
| Fig. 3, SI Video 3 | Whole body of a mouse pup (P14), CX3CR1 line. | Propidium iodide staining (561 excitation), nanobooster Atto647 for immune cells (microglia cells in mouse brain)<br>vDISCO whole-body staining | 561 nm,<br>Scanning:<br>f_galvo = 49.99 Hz<br>z-step = 15 μm<br>galvo_amp = 3.0 V | PlatinumTL™ 0.9x/0.045 telecentric lens | 6 tiles/ch, dual illumination, 233 GB. Active (fan) cooling of galvos to avoid overheating.<br>Imaging only with the moving cuvette (without the stationary cuvette), filed with DBE, by using focus tracking and ETL parameter adjustment depending on cuvette position. |
| Fig. 3, SI Video 4 | Chicken embryo with neurofilament staining at E9 (#4543) | Neurofilament staining with M Anti-RMO270, G Anti-Mouse Cy3<br>BABB clearing | 488 nm (auto-fluorescence), 561 nm (neurofilament)<br>z_step = 5 μm | Lensagon 1.2x/0.08 telecentric lens | 2 tile/ch, 230 GB total. |
| Fig. 3, SI Video 5 | Mouse brain, APPPS-1 mouse | Staining: hFTAA (amyloid plaques)<br>SMA-Cy3 (arterial vessels)<br>iDISCO clearing | 488 nm (hFTAA), 561 nm (SMA-Cy3)<br>z_step = 3 μm | Lensagon 1.2x/0.08 telecentric lens | 1 tile/ch, 136 GB total. |
| | Mouse brain, Vglut2-Cre mouse | Retrograde AAV cre-dependent variant of td-Tomato (ssAAV-retro/2-CAG-dlox-tdTomato (rev)-dlox-WPRE - bGHp(A)<br>iDISCO clearing | 561 nm<br>z_step = 2 μm | Olympus XLFluor-340, 4x/0.28 | 1 tile/ch, 70 GB total. |
| Fig. 3, SI Video 6 |  |  |  |  |  |
| SI Video 7 | Mouse brain, hippocampus region | Thy1-YFP AF-561<br>iDISCO clearing | 561 nm<br>z_step = 2 μm | Mitutoyo BD Plan Apo 5X/0.14 | 1 tile/ch, 65 GB total. Sample immersed in moving cuvette (5x10x45 mm) filled with DBE, dipped into a stationary chamber (40x40x50 mm) filled with DBE. |
| Fig. 3, SI Video 8 | Human cortex, occipital lobe 2 | Staining Neutral Red.<br>Clearing with MASH-NR (ECi) protocol. | 561 nm<br>z_step = 5 μm | Mitutoyo BD Plan Apo 5X/0.14 | 6 tiles/ch, 129 GB total. Sample immersed in dipping cuvette (10x20x45 mm) filled with ECi, dipped into a stationary chamber (40x40x75 mm) filled with DBE. |
| SI Video 9 | Human cortex, occipital lobe 1 | Staining with Methylene Blue/<br>Clearing with MASH-MB (ECi) protocol. | 638 nm<br>z_step = 5 μm | Mitutoyo BD Plan Apo 5X/0.14 | 12 tiles/ch, 110 GB total, clamped and dipped into a stationary chamber (40x40x50 mm) filled with ECi. |
| SI Video 10. | Xenopus tadpole stage 58 (44 days post-fertilization) | Primary AB: Atp1a1,<br>secondary: Alexa-Fluor-555 goat anti-mouse IgG.<br>BABB clearing | 561 nm<br>z_step = 7 μm | Mitutoyo BD Plan Apo 5X/0.14 | 40 tiles/ch, 1.68 TB, sample immersed in moving cuvette (10x10x45 mm), dipped in stationary cuvette (40x40x75 mm), both filled with DBE. |

For all datasets, stitching and fusion was performed in Fiji/BigStitcher, visualization in Imaris.
\* Detection filter: QuadLine Rejectionband ZET405/488/561/640.
\*\* Default scanning parameters: $t_{exp}$ = 20 ms, $f_{galvo}$ = 99.99 Hz, $galvo_{amp}$ up to 3 V depending on FOV size.

**SI Table 4.** sCMOS cameras supported by the mesoSPIM control software.

| Model | FOV: W x H, mm | Sensor diagonal, mm | Pixel array | Pixel size, $\mu\text{m}$ | Mega-pixels | Status | Notes | mesoSPIM version |
| --- | --- | --- | --- | --- | --- | --- | --- | --- |
| Hamamatsu Orca 4.0v3 | 13.312 x 13.312 | 18.8 | 2048 x 2048 | 6.5 | 4.2 MP | Supported and recommended | Use the version with PCIe frame grabber, not the USB3 (the latter is too slow) | v4-5 |
| Hamamatsu Orca Fusion | 14.976 x 14.976 | 21.2 | 2304 x 2304 | 6.5 | 5.3 MP | Tested, stable | Use CoaXPress frame grabber version, NOT the USB3. | v4-5 |
| Hamamatsu Orca Lightning | 25.344 x 14.256 | **29.1** | 4608 x 2592 | 5.5 | 12 MP | Tested, stable | High resolution across entire FOV only with large-FOV detection objectives. | V4-5 upgrade option |
| Teledyne Photometrix Kinetix | 20.8 x 20.8 | **29.4** | 3200 x 3200 | 6.5 | 10.2 MP | Tested, stable | High resolution across entire FOV only with large-FOV detection objectives. | v4-5 |
| Teledyne Photometrix Prime BSI | 13.312 x 13.312 | 18.8 | 2048 x 2048 | 6.5 | 4.2 MP | Tested, use at your own risk |  | v4-5 |
| Teledyne Photometrix Iris 15 | 21.49 x 12.61 | **24.9** | 5056 x 2960 | 4.25 | 15 MP | Supported and recommended | High resolution across entire FOV only with large-FOV detection objectives. | Benchtop and v4-5 upgrade |

**SI Table 5.** Stages and motion controllers supported by the mesoSPIM.

| Controller | Stages | Notes | mesoSPIM version |
| --- | --- | --- | --- |
| PI C-884 | L-509.20DG10, L-509.40DG10, M-060.DG, M-406.4PD | Serves up to 6 stages, including rotation. | v4-5 default controller |
| PI C-663 | Any of the PI stages above | Serves 1 stage, multiple controllers can be used. | v5 option |
| ASI Tiger (TG8-BASIC) | LS-50, LS-100, C60-3060 | Serves 6 motors with 3 additional free slots. | Benchtop |

Abbreviations: PI, Physik Instrumente; ASI, Applied Scientific Imaging.

**SI Table 6.** Filter wheels supported by the mesoSPIM.

| Model | Max. filter diameter, mm | N(filter slots) | Notes | mesoSPIM versions |
| --- | --- | --- | --- | --- |
| ZWO EFW-MINI | 31 mm | 5 | Very compact and low-cost astronomy filter wheel, with integrated controller and USB cable. | Benchtop |
| **Ludl**<br>**96A350** | 32 mm | 10 | Large filter wheel with separate controller (MAC6000), serial cable, and high cost. | v4-5 |
| **Sutter**<br>**Lambda 10** | 25 | 10 | deprecated | Used in early versions. |

**SI Table 7.** Parts list for mesoSPIM v4-5 upgrade.

All prices are approximate and subject to change at any time by the vendors.
| Part | Qty | Cost/unit, USD |
| --- | --- | --- |
| Mitutoyo M Plan Apo 2x/0.055 | 1 | 1,100 |
| Mitutoyo M Plan Apo 5x/0.14 | 1 | 900 |
| Mitutoyo M Plan Apo 7.5x/0.21 | 1 | 1,800 |
| Mitutoyo M Plan Apo 10x/0.28 | 1 | 1,200 |
| Mitutoyo G Plan Apo 20x(t3,5) | 1 | 3,600 |
| Mitutoyo MT-1 tube lens | 1 | 900 |
| Mitutoyo 378-726D motorized turret (revolver) | 1 | 4,600 |
| Photometrics Iris 15 camera | 1 | 12,000 |
| Other hardware | - | 800 |
| Total, USD |  | 26,900 |

The detection filters as well as focusing and sample stages can be re-used.

**SI Table 8.** Estimated acquisition time and file size for imaging 1 cm^3^ sample on the Benchtop-mesoSPIM.

| Magnification | Fast mode (z-step 5 μm) | | Nyquist mode (z-step 2 μm) | |
| --- | --- | --- | --- | --- |
|  | Minutes, per 1 cm<sup>3</sup> actual**, average by vol. | File size, GB actual, average by vol. | Minutes, per 1 cm<sup>3</sup> actual, average by vol. | File size, GB actual, average by vol. |
| 1x | 13, 6 | 59, 27 | 33, 15 | 149, 68 |
| 2x | 53, 24 | 239, 109 | 133, 60 | 598, 272 |
| 5x* | 200, 152 | 897, 681 | 500, 380 | 2244, 1704 |
| 7.5x | 373, 342 | 1'676, 1534 | 933, 854 | 4190, 3835 |
| 10x | 720, 607 | 3232, 2727 | 1800, 1518 | 8081, 6818 |
| 20x | 2640, 2430 | 11'852, 10908 | 6600, 6074 | 29632, 27272 |

\* For comparison, imaging at this magnification with mesoSPIM v.5 (camera Hamamatsu Orca Flash 4.3) in fast mode takes 333 instead of 200 min and produces 419 instead of 897 GB of data.
\*\* Actual time is larger than (volume) average time because the the number of tiles in actual acquisition is always integer, which may be non-optimal for small volumes. At high magnifications and/or large volumes the difference between the two estimates becomes negligible.

**SI Table 9:** Simulated excitation beam waist diameter (FWHM) as a function of imaging medium thickness.

| "DBE" thickness | Beam waist FWHM (cross-section), $\mu\text{m}$ | Spherical aberration and defocus (3-rd order) Zernike term, waves | Total wavefront RMS, waves |
| --- | --- | --- | --- |
| **0 mm (air)** | **1.8 $\mu\text{m}$** | **0** | **0** |
| **5 mm** | **1.9 $\mu\text{m}$** | **-0.04 $\lambda$** | **0.07 $\lambda$** |
| **10 mm** | **2.4 $\mu\text{m}$** | **-0.08 $\lambda$** | **0.14 $\lambda$** |
| **15 mm** | **3.2 $\mu\text{m}$** | **-0.12 $\lambda$** | **0.21 $\lambda$** |
| 20 mm | 3.6 $\mu\text{m}$ | -0.16 $\lambda$ | 0.29 $\lambda$ |
| 30 mm | 3.0 $\mu\text{m}$ | -0.23 $\lambda$ | 0.43 $\lambda$ |
| 40 mm | 4.0 $\mu\text{m}$ | -0.31 $\lambda$ | 0.57 $\lambda$ |

**SI Table 10:**
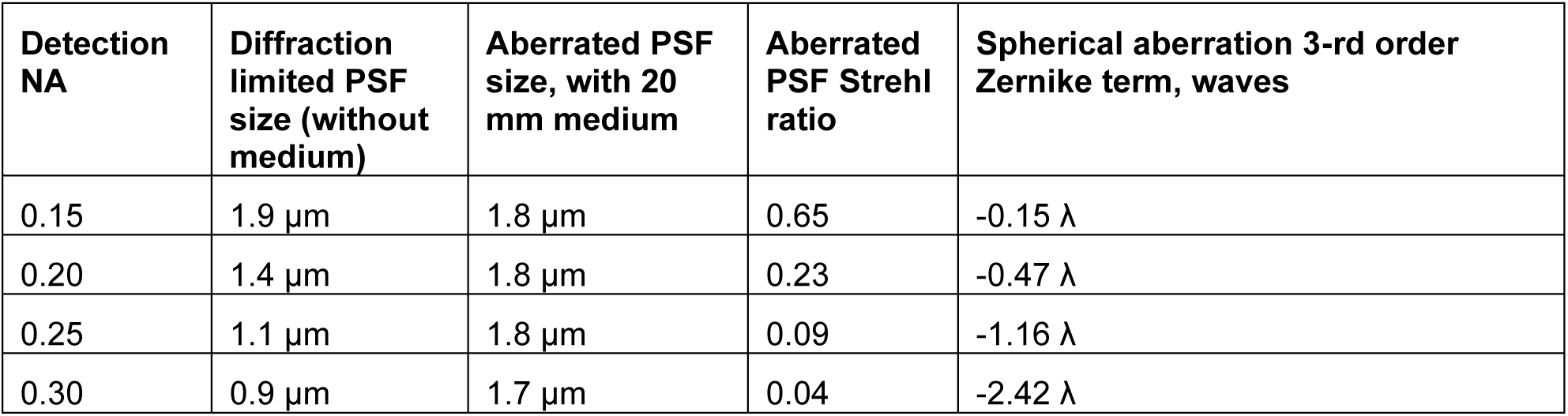
Simulated detection performance as a function of detection NA.

| Detection NA | Diffraction limited PSF size (without medium) | Aberrated PSF size, with 20 mm medium | Aberrated PSF Strehl ratio | Spherical aberration 3-rd order Zernike term, waves |
| --- | --- | --- | --- | --- |
| 0.15 | 1.9 $\mu\text{m}$ | 1.8 $\mu\text{m}$ | 0.65 | -0.15 $\lambda$ |
| 0.20 | 1.4 $\mu\text{m}$ | 1.8 $\mu\text{m}$ | 0.23 | -0.47 $\lambda$ |
| 0.25 | 1.1 $\mu\text{m}$ | 1.8 $\mu\text{m}$ | 0.09 | -1.16 $\lambda$ |
| 0.30 | 0.9 $\mu\text{m}$ | 1.7 $\mu\text{m}$ | 0.04 | -2.42 $\lambda$ |

**SI Table 11:** Simulated detection performance as a function of medium refractive index.

Detection NA=0.15, medium thickness 20 mm.
| Medium R.I. | Aberrated PSF size | Aberrated PSF Strehl ratio |
| --- | --- | --- |
| 1.33 | 1.8 $\mu\text{m}$ | 0.72 |
| 1.47 | 1.8 $\mu\text{m}$ | 0.66 |
| 1.52 | 1.8 $\mu\text{m}$ | 0.65 |
| 1.56 | 1.8 $\mu\text{m}$ | 0.65 |
| 1.60 | 1.8 $\mu\text{m}$ | 0.64 |
| 1.80 | 1.8 $\mu\text{m}$ | 0.64 |
| 1.99 | 1.8 $\mu\text{m}$ | 0.65 |

**SI Table 12:** Simulated detection performance as a function of medium thickness.

Detection NA=0.15, medium R.I.=1.56. Diffraction-limited cases (Strehl ratio $>\sim 0.80$ ) are marked in **bold**.
| Medium thickness | Aberrated PSF size | Aberrated PSF Strehl ratio |
| --- | --- | --- |
| **5 mm** | **1.9 $\mu\text{m}$** | **0.97** |
| **10 mm** | **1.8 $\mu\text{m}$** | **0.89** |
| **15 mm** | **1.8 $\mu\text{m}$** | **0.78** |
| 20 mm | 1.8 $\mu\text{m}$ | 0.65 |
| 30 mm | 1.6 $\mu\text{m}$ | 0.40 |

